# Defining a Nonribosomal Specificity Code for Design

**DOI:** 10.1101/2022.08.30.505883

**Authors:** Aleksa Stanišić, Carl-Magnus Svensson, Ulrich Ettelt, Hajo Kries

**Affiliations:** Independent Junior Research Group Biosynthetic Design of Natural Products, Leibniz Institute for Natural Product Research and Infection Biology e.V., Hans Knöll Institute (HKI Jena), Beutenbergstr. 11a, 07745 Jena, Germany; Applied Systems Biology, Leibniz Institute for Natural Product Research and Infection Biology e.V., Hans Knöll Institute (HKI Jena), Beutenbergstr. 11a, 07745 Jena, Germany

**Keywords:** nonribosomal peptide synthetase, enzyme engineering, adenylation, multispecificity, surfactin

## Abstract

Nonribosomal peptide synthetases (NRPSs) assemble bioactive peptides from an enormous repertoire of building blocks. How binding pocket residues of the nonribosomal adenylation domain, the so-called specificity code, determine which building block becomes incorporated has been a landmark discovery in NRPS enzymology. While specificity codes enable the prediction of substrate specificity from protein sequence, design strategies based on rewriting the specificity code have been limited in scope. An important reason for failed NRPS design has been that multispecificity has not been considered, for a lack of suitable assay formats. Here, we employ a multiplexed hydroxamate specificity assay (HAMA) to determine substrate profiles for mutant libraries of A-domain in the termination module the SrfAC of surfactin synthetase. A generalist version of SrfAC is developed and the functional flexibility of the adenylation reaction is probed by fully randomizing 15 residues in and around the active site. We identify mutations with profound impact on substrate selectivity and thus reveal a remarkable evolvability of A-domains. Statistical analysis of the specificity divergence caused by point mutations has determined the impact of each code position on specificity, which will serve as a roadmap for NRPS engineering. The shortness of evolutionary pathways between NRPS specificities explains the rich natural substrate scope and suggests directed evolution guided by A-domain promiscuity as a promising strategy.

## Introduction

Promiscuous enzyme activities serve as evolutionary springboard towards novel functions (1–3). It is believed that in natural evolution, multispecific, generalist ancestor enzymes have gained specificity under strong selective pressure. Directed evolution imitates this process in the laboratory to design customized enzymes with broad applications (4–7). To generate suitable starting points for directed evolution, specialized enzymes can be reverted to promiscuous, ancestor-like states by amplifying weak activities towards noncognate substrates (8). These generalist enzymes serve as a starting point for re-specialization towards a desired function in laboratory evolution (9). Not all enzymes are equally evolvable, however. It has been shown that enzyme families with high natural functional diversity are more amenable to change than those fulfilling identical roles across the homology tree (10). Secondary metabolism is especially enriched with promiscuous activities resulting in a mishmash of natural product congeners (11– 13). Hence, enzymes from secondary metabolism seem especially suitable for studying promiscuity and mapping evolutionary pathways between different functions (14).

Nonribosomal peptides (NRPs) are a class of secondary metabolites of great importance for human use as antibiotics, immunosuppressants, and anti-cancer drugs (15). NRPs are assembled on large multidomain enzymes termed nonribosomal peptide synthetases (NRPSs) (16). NRPSs consist of enzyme domains catalysing individual reactions. Domains are grouped in modules, each of which incorporates a single substrate into the peptide chain in an assembly line fashion (Figure 1a). Substrates are first activated with ATP by adenylation (A-)domains before being tethered to thiolation (T-)domains and condensed with the substrate from the adjacent module by condensation (C-)domains. The release of the final product is typically achieved by a terminal thioesterase (TE-)domain catalysing hydrolysis or intramolecular cyclization of mature linear peptide. The large variety of NRPS architectures and corresponding NRP products must result from fast evolutionary diversification. Sequences of NRPS gene clusters suggest evolutionary mechanisms relying on a combination of gene recombination and neofunctionalization (17–22). However, the neofunctionalization mechanisms of individual modules have remained largely elusive because nonribosomal multispecificity has been cumbersome to analyze (23, 24).

**Figure 1.**
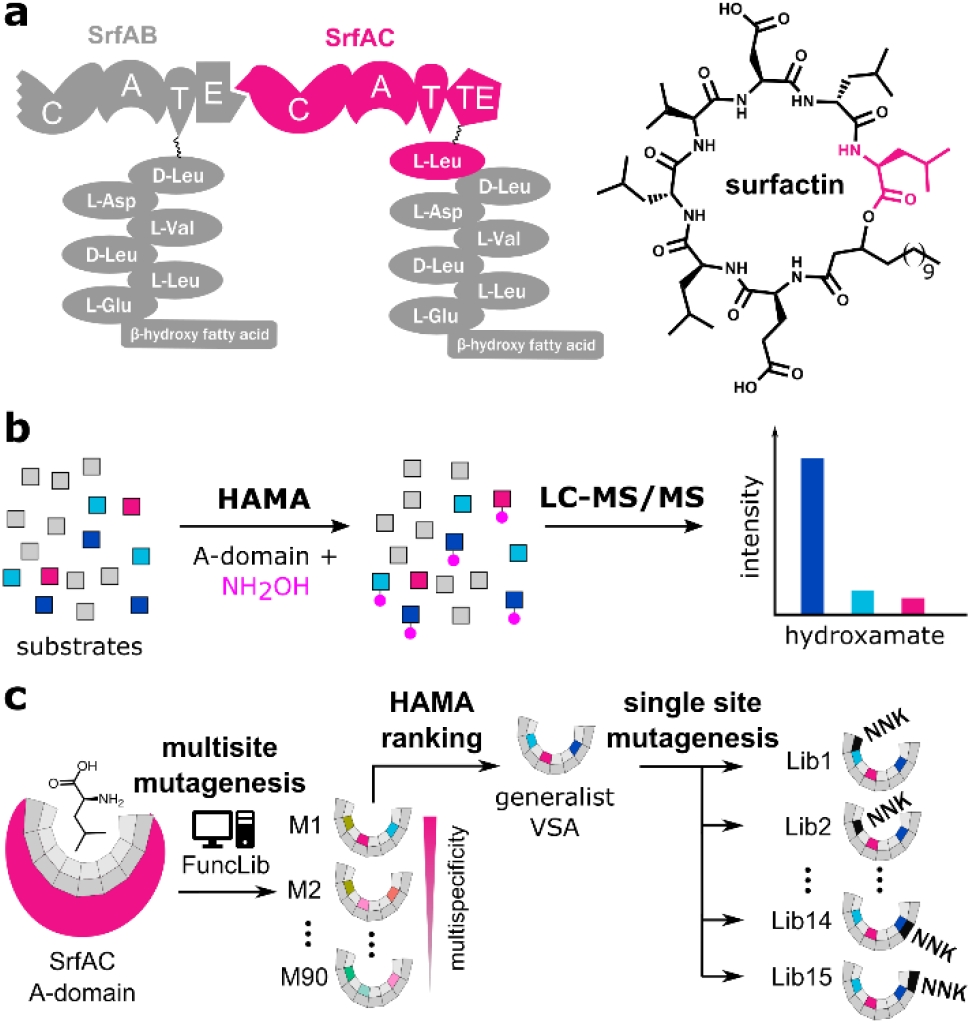
a) The nonribosomal peptide synthetase module SrfAC incorporates the terminal leucine into surfactin. b) The hydroxamate assay (HAMA) records substrate profiles of nonribosomal adenylation (A-)domains (38). c) The substrate profile of Leu-specific SrfAC is diversified by multisite mutagenesis aided by the FuncLib algorithm (M1: mutant 1) (8), followed by site-directed saturation mutagenesis at 15 positions close to the A-domain active site (Lib1: library 1) using degenerate NNK codons.

The modular nature of NRPSs makes them an attractive engineering target for sourcing custom-made peptides. Controlling A-domains, which in turn control the identity of activated and incorporated substrates, would unlock efficient biosynthetic drug design (25). Natural A-domains recognize more than 500 different monomers (26, 27) and can be highly specific (28), bispecific (29), or multispecific (14, 19). Structural and sequence analyses have revealed conservation of ‘specificity code’ residues in the binding pockets of A-domains activating the same substrate (30– 32). The initial 8-residue code, later amended by 2^nd^ and 3^rd^ shell residues, allowed the development of algorithms predicting the identity of the final products from NRPS protein sequence (33–35).

When NRPS reprogramming was attempted based on A-domain specificity codes, successful switches were limited to structurally similar substrates indicating that specificity signatures are not readily transferable between A-domains. Nevertheless, good designability of A-domain specificity has been demonstrated on Phe-specific GrsA which acquired a 5×10^5^-fold switch in specificity towards “click” amino acid propargyl-Tyr by introducing a single mutation in the binding pocket (24). High-throughput screening using yeast surface display has further bypassed the limitations of rational A-domain design (36, 37), but this approach remains limited to substrates with a covalent binding handle. For efficient A-domain engineering, it is essential to better understand how binding pocket mutagenesis impacts specificity profiles. However, straightforward adenylation assays for A-domain multispecificity have been lacking. In previous work, we have developed a hydroxamate assay (HAMA, Figure 1b) to determine a complete specificity profile of an A-domain in a single reaction, dramatically reducing the workload and facilitating the determination of A-domain multispecificity (38).

Here, we take advantage of HAMA to investigate the impact of mutations on the specificity landscape of the A-domain from SrfAC, the termination module from surfactin synthetase in *B. subtilis*. SrfAC is a standalone module with C-A-T-TE architecture incorporating the terminal L-Leu into the biosurfactant surfactin (Figure 1a). We harness HAMA to unravel the evolutionary pathways leading towards diverse specificities via an intermediate with broadened substrate spectrum (Figure 1c). Computationally-supported introduction of multisite mutations into module SrfAC has yielded variant VSA with enhanced activity towards several substrates and retained stability. In a library of single mutations prepared from VSA, we demonstrate high flexibility of adenylation specificity. Quantitative understanding of the mutational landscape of NRPS specificity and a refined specificity code will serve as a roadmap for the future engineering of novel bioactive molecules.

## Results

### Converting SrfAC into a generalist

Assuming that promiscuous activities are evolutionary stepping stones towards novel functions, our first aim was to develop a variant of SrfAC with relaxed specificity (Figure 2). Module SrfAC incorporates the terminal L-Leu residue of surfactin (Figure 1a), has been thoroughly investigated with biophysical methods (39, 40), and shows stable expression in *E. coli* even in microtiter plate format (Figure S1). Interestingly, SrfAC shows substantial activity towards nonproteinogenic D-Val, which is presumably inconsequential to the host *B. subtilis* due to the absence of this stereoisomer in the organism. Aside from D-Val, SrfAC shows low, but detectable side activities that are ca. 20-fold (l-Ile, l-Cys, l-Val, l-Met) or 340-fold (L-Phe) lower than the main activity for L-Leu (Figure 2c). In nature, surfactins are produced as a mixture of congeners, including [Val7]surfactin (41), which presumably results from the weak side-activity of the SrfAC A-domain. Good protein behaviour combined with weak natural multispecificity make this system ideal for studying NRPS promiscuity (42).

**Figure 2.**
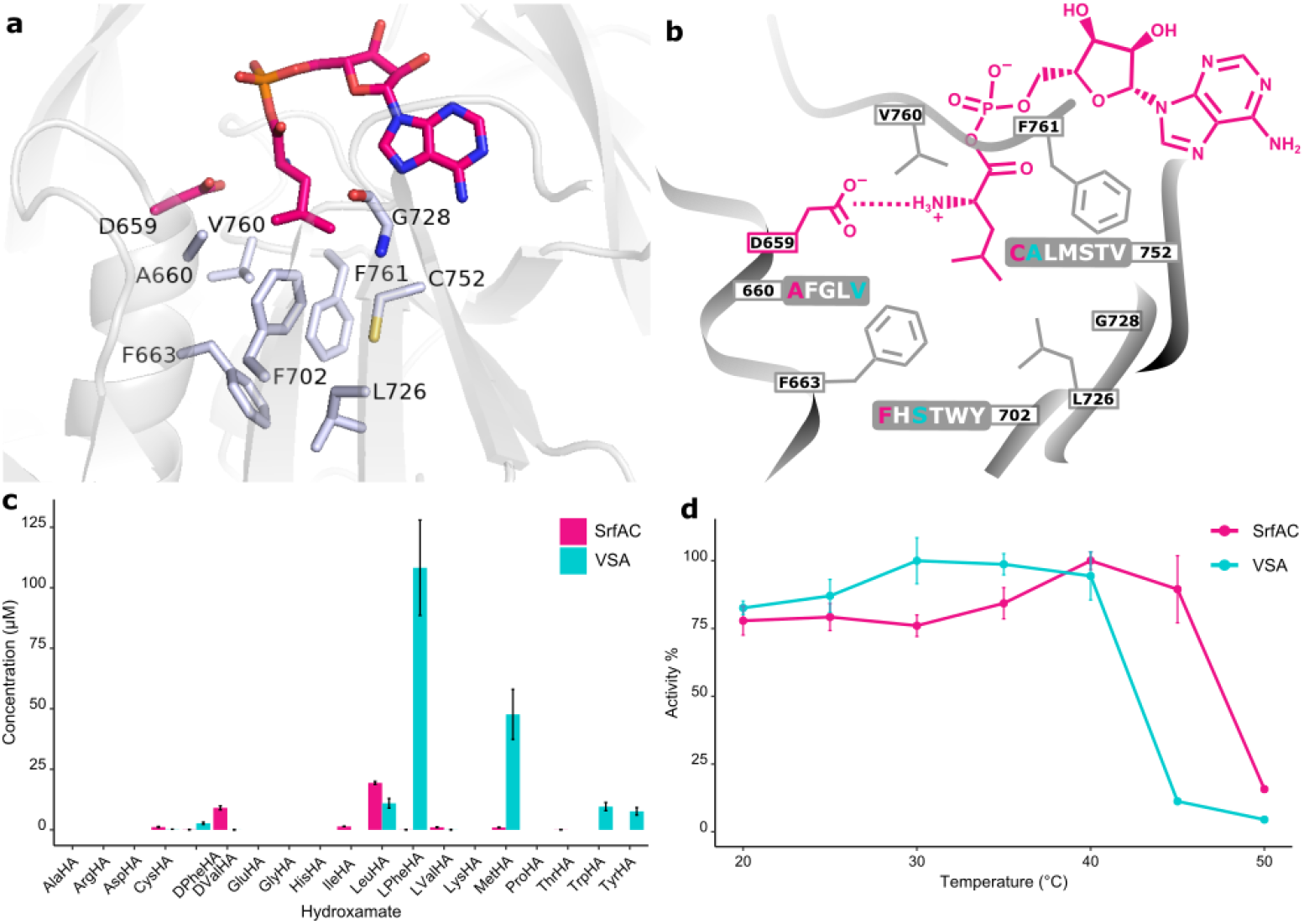
SrfAC and mutant VSA with relaxed specificity. a) Binding pocket of SrfAC homology modelled with Yasara(54) in complex with Leu-AMP (sticks in pink). Specificity code residues are shown as grey sticks. The model is built against the SrfAC crystal structure as a template (PDB:2VSQ).(39) b) Mutagenesis of the SrfAC binding pocket. Three specificity code residues of SrfAC selected for randomization (660, 752, and 702) and the list of tolerated residues predicted by FuncLib (8) are marked. SrfAC and VSA residues are labelled in pink and cyan, respectively. c) HAMA specificity profiles. Enzyme reactions were incubated for 60 min at 25 °C with 1 µM enzyme and a mixture of amino acids. d) Thermostability. Enzyme reactions were incubated for 20 min at different temperatures and 1 µM enzyme and the production of hydroxamates was followed. Error bars are standard deviations from three c) or two d) technical replicates from two batches of enzyme.

Multisite mutations can functionally diversify enzyme active sites but have a high risk to be deleterious for activity and stability. To gain a functional repertoire of diverse SrfAC multisite mutants, we took advantage of FuncLib, an automated algorithm using phylogenetic analysis and Rosetta modelling to predict the tolerability of mutations (8). FuncLib filters out mutations likely to result in inactive variants. To do so, FuncLib scores single site mutants present in a multiple sequence alignment using the Rosetta algorithm, which offers a good approximation of protein stability (43). For use of the FuncLib webserver, a model of SrfAC in complex with Leu-AMP was built using the YASARA modelling software (Figure S2) (44). The Leu-AMP ligand and the invariable catalytic residue D659 (Figure 2a) were fixed during the calculations. From the eight positions of the specificity code (A660, F663, F702, L726, G728, C752, V760, F761), we selected three for experimental multisite randomization based on experience from previous A-domain engineering campaigns (24, 36). Being located at the entrance (A660 and C752) and the bottom (F702) of the binding pocket, a decisive impact on substrate recognition was expected. At these three positions, the most beneficial 5, 6, and 7 residue identities according to the first stage of the FuncLib protocol were combined in a random library (Figure 2b). The library of 210 triple mutants was cloned by combining oligonucleotides bearing degenerate codons for each position in appropriate ratios (Table S1 and 2). As expected for an A-domain activating a nonpolar amino acid, FuncLib predicted mutational tolerance towards residues with predominantly aliphatic or hydrophobic side chains (Figure 2b).

To analyze the triple mutants, protein was expressed in four 96-well microtiter plates from randomly picked colonies and specificity profiles with 17 proteinogenic and two nonproteinogenic substrates were measured with HAMA. The strength of the FuncLib prediction is illustrated by 46% of library members having detectable activity – remarkably high considering significant losses in activity typically accompanying multisite A-domain mutagenesis. Out of the candidates with highest activity and promiscuity, triple mutant A660<u>V</u>-F702<u>S</u>-C752<u>A</u> (VSA) was selected for further characterization. VSA shows enhanced activity for several substrates and activates Phe and Met in HAMA at rates surpassing SrfAC with Leu. Additionally, activity of VSA for Leu, Trp, and Tyr is high (Figure 2c). The large-to-small mutation F702S presumably frees up space for bulkier, hydrophobic amino acid substrates such as Phe and Met. The enhanced turnover for multiple substrates predisposes VSA for further diversification because the high activity lifts several substrates above the detection threshold.

In addition to broad substrate tolerance, an ancestor-like enzyme must be stable enough to withstand further mutations (45). To test thermal stability, the adenylation activity of VSA was followed at a range of temperatures between 20 and 50 °C (Figure 2d). VSA maintains full activity up to 40 °C, indicating the absence of major stability trade-offs in comparison with the parent SrfAC. To characterize the effect of the VSA mutations, saturation kinetics with the three major substrates (L-Leu, L-Phe and L-Met) were measured with the MesG/hydroxylamine assay (Figure S3). The adenylation rate *k*_cat_ for all three substrates is maintained at wild type-like levels with differences originating in *K*_M_ values. *K*_M_(L-Leu) shows a 50-fold increase from 10 µM in SrfAC to 500 µM in VSA while *K*_M_(L-Phe) and *K*_M_(L-Met) of VSA are within 2 and 10-fold of *K*_M_(L-Leu). Consequently, specificity constants *k*_cat_/*K*_M_ of all three substrates fall within one order of magnitude. Combining good stability and an expanded substrate repertoire at wild type-like rates, VSA was ideally suited for further functional diversification.

### Broad functional sequence space in VSA mutants

Having established VSA as a robust mutant with broad specificity, we proceeded to thoroughly probe the effects of single point mutations on the specificity profile of VSA. We exhaustively covered the binding pocket with site-saturation mutagenesis libraries in 15 positions (Figure 3a, Figure S4). In addition to the 8 specificity code residues in the first shell, 7 second shell residues were included. To maintain 90 % coverage of each NNK library, we screened 92 colonies per library with HAMA in microtiter plates. Libraries were sequenced and completed by individually cloning mutants missing from them (Table S5). Activity was detected in 50 % (147/300) of mutants from all libraries. Out of 19 offered substrates, 11 yielded detectable products with at least some of the mutants (Figure 3b, Figure S5). The set of active substrates was strongly biased towards low polarity but, interestingly, included D-configured Phe and Val (Figure S5).

**Figure 3.**
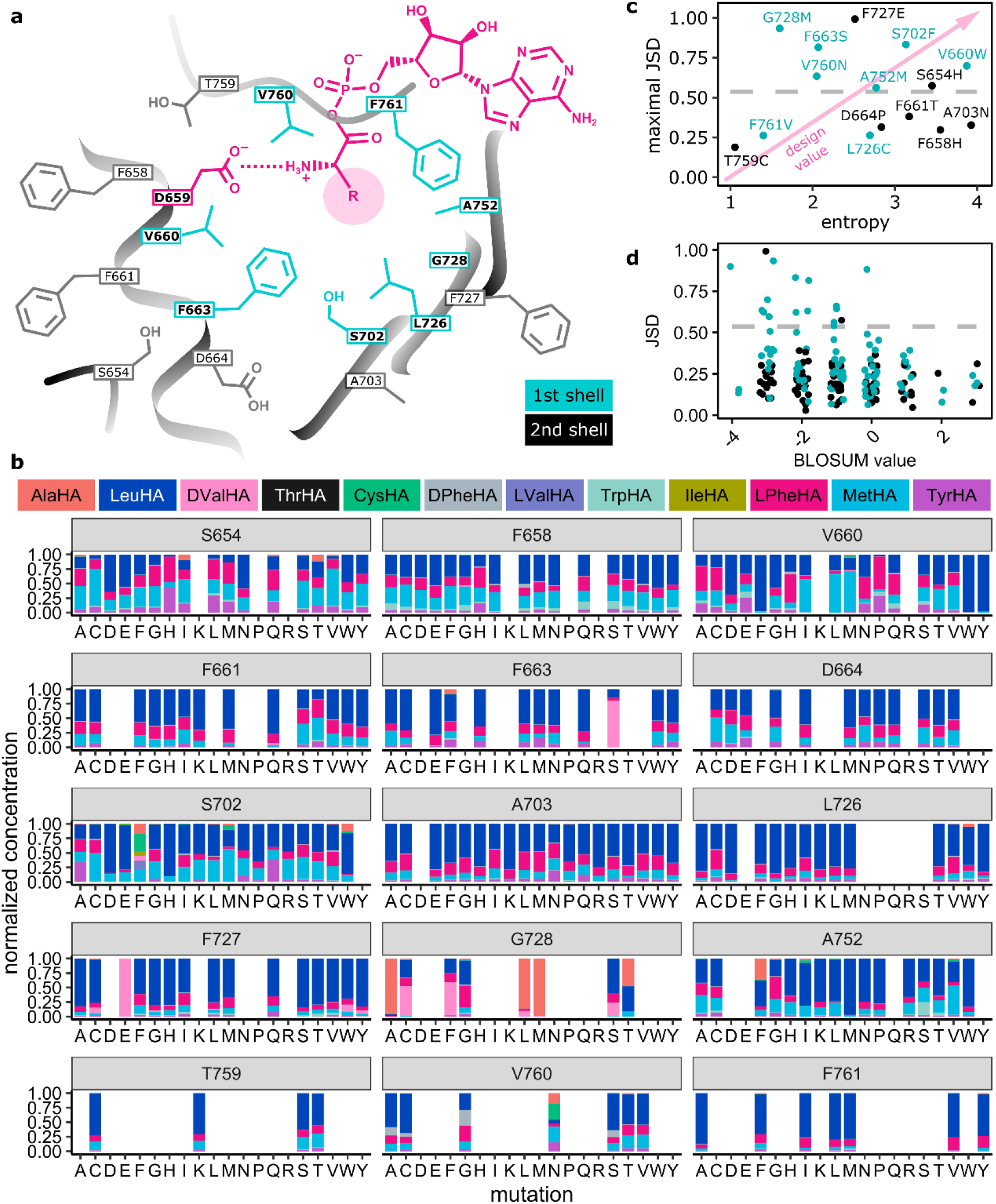
a) Scheme of the VSA binding pocket. b) HAMA specificity profiles from screening NNK libraries of VSA. When the sum of hydroxamates for one mutant is below 1% of the wild type, it is set to zero; all other mutants are normalized to one. Identical mutants have been averaged. c) Maximum Jensen-Shannon distance (JSD) per position (55) plotted against the entropy of the activity distribution at the same position (cyan: first shell, black: second shell). The dashed line indicates significant divergence from VSA. d) JSD for each mutant plotted against the BLOSUM score of the mutation, where more negative values indicate a larger mutational distance.

### Ranking positions for designability

By recording the specificity profiles of the A-domain mutants, we wanted to learn in which positions mutations caused the largest specificity changes. Therefore, we ranked all 15 positions according to the largest difference in specificity profile obtained with any mutant at that position (Figure 3c). The difference in specificity was quantified as the Jenson-Shannon distance (JSD) of the hydroxamate distributions. Mutants with less than 1% of the wild type activity were excluded from the analysis to avoid artefacts. Interestingly, one of the furthest distances from the wild type was observed with an F663S mutation at the floor of the binding pocket, which is strikingly similar to a Trp to Ser mutation that earlier showed a powerful effect on substrate specificity of GrsA (24). As expected, the maximum JSD was generally higher for first than for second shell residues. Notable exceptions are the second shell mutations F727E and S654H showing large divergences and first shell positions F761 and L726 remaining uninfluential.

In addition to the maximum divergence of specificity, the activity level of mutants is an important parameter to gauge the usefulness of a position for A-domain design. Activity distributions for all mutants in one position were represented as entropy values (Figure 3c), where a high entropy value indicates a uniform distribution of activity and therefore high mutational tolerance. Low entropy values indicate sensitive positions where all mutations strongly reduce activity compared to the wild type. The most useful positions for design will combine a large divergence of the specificity profile with robust activity (top right corner of Figure 3c). For instance, first shell positions 702 in the beta-sheet and 660 adjacent to the conserved D659 have been identified as particularly valuable for design, because they give rise to diverse specificity at good activity levels. At the other extreme, at position T759, activity is almost destroyed only by a conservative mutation to Ser without yielding a noticeable difference in specificity. As expected, specificity profiles generally diverged more strongly from the starting point when mutations were less conservative, as measured by the BLOSUM score (Figure 3d). Accordingly, polar (Asp, Glu, His, Lys, Arg), rigid (Pro) and bulky (Trp, Tyr) substitutions in the hydrophobic A-domain binding pocket create mostly inactive enzymes (Figure S6).

### Hit validation

Due to the large technical variability in our multistep microtiter plate screening protocol, we remeasured HAMA profiles for selected mutants after large-scale purification (Figure 4) to challenge the most remarkable results. Thus, it was confirmed that two mutants at position G728 (G728A, G728M) show high specificity for Ala, which is not a detectable substrate in SrfAC or VSA. Another mutant (V660W) achieves higher specificity for Leu than SrfAC by eliminating the side-activity for D-Val. Met is rarely encountered as NRPS building block but has been enhanced already in VSA compared to SrfAC. Several mutations in position 660 make Met the major substrate (e.g. V660I). Additionally, activation of D-configured Phe is favoured by Gly substitution at V760. The enhanced promiscuity of the S702F mutant observed in the screening experiment could not be corroborated in the validation experiment, where predominately Leu activity was detected (Figure S10).

**Figure 4.**
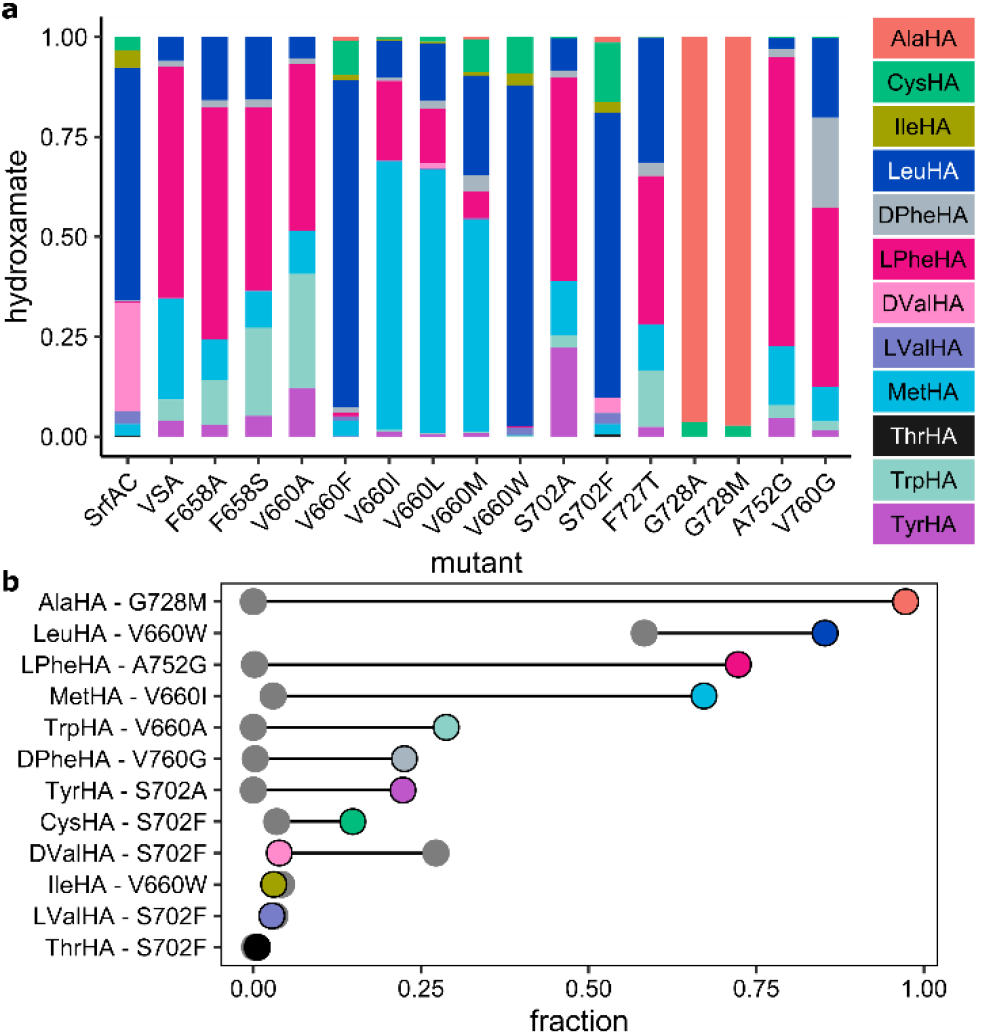
a) HAMA specificity profiles of selected VSA mutants. Activities are normalized to the highest hydroxamate peak for each mutant. Enzyme reactions were incubated for 60 min at 25 °C and 1 µM purified enzyme. Error bars show standard deviations from three technical replicates from two batches of enzyme. b) For each hydroxamate, the mutant showing the largest fraction in its product spectrum (colored circle) is compared to SrfAC (grey). For unnormalized activity levels, see Figure S11.

## Discussion

Decryption of the A-domain specificity code has been one of the most significant breakthroughs in NRPS enzymology (31, 32, 25). However, subsequent attempts to repurpose A-domains to produce tailored peptides were plagued with losses of activity (23, 46). While specificity signatures for individual substrates are well documented (33, 34), the rules and mechanisms governing (laboratory) evolution of substrate selection have remained elusive. A thorough study of A-domain promiscuity of tyrocidine synthetase TycA with a range of natural and synthetic substrates revealed a high specificity of TycA for Phe, with three orders of magnitude selectivity over the second best substrate Tyr (28). Laboratory evolution of TycA from Phe to Ala specificity yielded a moderately active enzyme and required several rounds of screening and mutagenesis. Only few projects succeeded to mutate A-domains inspired by specificity codes (46–48). Therefore, the question how natural evolution found pathways towards so many different substrates and how these pathways can be explored and extended by design is still open.

One main culprit for our poor understanding of A-domain neofunctionalization has been the lack of adequate specificity assays (25). With the development of HAMA profiling, this bottleneck has been cleared and a complete specificity profile under competition conditions is recorded within minutes (38). Here, we use HAMA adapted for microtiter plate screening to conduct an in-depth investigation of the A-domain multispecificity landscape. Following the paradigm that generalist enzymes are hubs of natural evolution (9, 1, 49, 3), we have developed a progenitor-like version of the NRPS module SrfAC as a stepping stone towards more diverse specificities. Compared to SrfAC, the triple mutant VSA shows remarkable 2000-and 50-fold increases for Phe and Met, respectively, and a generally higher level of hydroxamate formation with promiscuous substrates (Figure 2c).

While A-domain engineering typically focuses on one or few substrates, we aimed to expand into many directions by introducing single mutations into VSA and recording specificity profiles. The substrate range accessible at this small mutational distance from the generalist VSA proved remarkably diverse but not unlimited. While half of all single mutants were active for at least one of the aliphatic amino acids tested, no hydroxamates of polar amino acids were detected. Presumably, multiple binding pocket positions adapt to the general physico-chemical properties of the substrate, so that multiple residues must change to jump to a different class of substrate. Such an evolutionary jump may lead through a valley of low activity. Within one class of substrate, specificity switches are remarkably easy to achieve. With reference to SrfAC, full specificity switches (Ala, Phe, Met) or substantial improvements (Trp, D-Phe, Tyr, Cys) were confirmed for seven substrates (Figure 4b). Notably, our screening method has uncovered new specificity codes that do not exist in nature (Table S7 and Figure S8), for instance a code for Leu with Trp in position 660 (Figure 4). Codes for Leu have Ala in position 660 in 94% of the cases but never Trp. The potential for discovering novel codes for the same substrate has earlier been demonstrated by Throckmorten et al. (50).

Since specificity codes have been an unreliable blueprint for A-domain design, we aimed to better define which residues are useful for customizing specificity through random mutagenesis. To rank positions in the A-domain, we have considered both the divergence of specificity from wild-type and the sensitivity of activity to mutation. We applied these metrics on the single mutations introduced into VSA. The analysis revealed a good but not perfect agreement between the influential positions that we identified (Figure 3c) and the 8 canonical code positions in the first shell of the binding pocket. Two of the first shell positions (726, 761) showed no use for design because specificity profiles remained unaltered while activity suffered strongly. Very few mutations in the second shell yielded specificity profiles significantly different from the starting point. Therefore, by analysing even more distant residues, we would not expect fundamentally new insights for the designability ranking, even if finely tuned long-range interactions can be important to design a perfect catalyst (51). These insights will help to prioritise positions for the directed evolution of A-domain specificity in the future. Considering the plasticity of A-domain specificity upon single mutation, we predict that screening smart, medium-sized libraries, perhaps supported by artificial intelligence, will suffice to unlock various target substrates (52, 53).

## Conclusion

Here, we have utilized HAMA to conduct a thorough investigation of A-domain specificity after mutation. We demonstrate the strength of promiscuity-guided screening to create generalist A-domains as progenitors for further diversification. Introducing point mutations in generalist VSA at only a few positions has been sufficient to achieve large changes in specificity, often without impairment in activity. The specificity and activity profiles for 15 positions in the A-domain binding pocket region have allowed to rank these positions by their design value, which will serve as a roadmap for redesigning nonribosomal peptides. The ease by which single mutations switch the specificity of A-domains explains how evolution has been able to recruit so many building blocks for nonribosomal peptides.

## Acknowledgments

This work has been funded by the Deutsche Forschungsgemeinschaft (DFG, German Research Foundation) with Project ID 441781663. Furthermore, we acknowledge financial support by the Daimler und Benz Stiftung, the Fonds der Chemischen Industrie, and a Max-Buchner-Fellowship. We thank Prof. Donald Hilvert (ETH Zurich) for providing plasmid pTrc99a-srfAC.

## Notes

### Competing Interest Statement

The authors have declared no competing interest.

