## Supplementary Information for "Defining a Nonribosomal Specificity Code for Design"

### Protein models

The 3D-model of SrfAC in complex with an L-Leu-AMP ligand was built with YASARA (1). The model was aligned to the crystal structure of SrfAC (PDB: 2vsq) to confirm that modelling did not change the position of specificity code residues in the binding pocket (Figure S2). The 3D-model of the VSA mutant was created on the SWISS-MODEL server (https://swissmodel.expasy.org/) (2, 3) by using the crystal structure of SrfAC (PDB: 2vsq) (4) in its thiolation state as a template. Both models were then aligned in PyMOL (https://pymol.org/).

### Cloning

#### *General cloning*

For the transformation of In-Fusion cloning reactions, *E. coli* HST08 Stellar competent cells were used (Takara Bio Europe). For the propagation and storage of plasmids, *E. coli* NEB 5-alpha (New England Biolabs) was used. Plasmid DNA, DNA fragments, and PCR products were purified using NucleoSpin kits (Macherey Nagel). DNA amplification was done with Q5 polymerase (New England Biolabs) following the supplier's instructions. Two-fragment cloning in linearized vector was done using the InFusion cloning kit (Takara Bio Europe). Oligonucleotide primers were made by custom synthesis (Eurofins Genomics) and sequence confirmation of assembled constructs was performed using the Mix2Seq service for Sanger sequencing. The plasmid pTrc99a-srfAC was kindly provided by Prof. Donald Hilvert (ETH Zurich). For the linearization of pTrc99a-srfAC, restriction enzymes BlpI, DraIII and BstBI (New England Biolabs, Massachusetts) were used, depending on the position of the mutation. Libraries were generated by amplification of one or two DNA fragments with primers bearing randomized codons. If the insert was amplified in two fragments, these were concatenated by assembly PCR to increase cloning efficiency. Cloned libraries were transformed into *E. coli* HST08 Stellar competent cells. Libraries were purified from overnight liquid cultures grown under ampicillin selection and resulting plasmid DNA was used to transform the protein expression strain *E. coli* HM0079 (5).

#### *Sequencing*

The identity of purified plasmids was confirmed by Sanger sequencing (Eurofins Genomics). The identity of library mutants was determined by Sanger sequencing of 96-well plates (Microsynth) containing aliquots taken from the saturated preculture shortly before induction of protein expression.

### Library design

#### *FuncLib library of SrfAC*

*SrfAC randomization with FuncLib*

To identify the essential residues required for adenylation, a SWISS homology model of the SrfAC A-domain was built on the crystal structure of EntF (PDB: 5T3D) solved in complex with serine adenosine vinylsulfonamide inhibitor (Ser-AVS), a nonhydrolyzable analogue of serine-AMP. A structure of L-Leu-AMP was modelled into the SrfAC homology model using YASARA. The resulting model was used for FuncLib randomization (6). Informed by the structural model, eight specificity code residues of SrfAC were selected for simultaneous randomization by FuncLib: A660, F663, F702, L726, G728, C752, V760 and F761 at default parameters for multiple sequence alignments (Min ID: 35, Max targets: 4000, Coverage: 75, E value: 0.0001). Conformations of AMP and D659 were fixed to maintain the interactions necessary for adenylation. FuncLib generated signatures of residues tolerated at each of the 8 selected positions. Three residues were selected (in bold) for the following construction of a library of triple mutants.

| | | | |
|---|---|---|---|
| **A660: AFGLV** | F663: FIMWY | **F702: FHSTWY** | L726: LIMVY |
| G728: GACS | **C752: CALMSTV** | V760: VACFILTY | F761: FHILMWY |

*Cloning of FuncLib library of SrfAC*

For generating a library of SrfAC triple mutants inspired by FuncLib, a series of oligonucleotides containing degenerate codons coding for predicted residues at the selected positions 660, 702, and 752 were used. The wild type residue was included in each position. For each randomized position, degenerate oligonucleotides were combined in ratios reflecting the number of possible codons and used for PCR amplification of DNA fragments with pTrc99a-SrfAC as a template (Table S1). A single DNA fragment for each position was generated resulting in three fragments (A, B, C). These were assembled by PCR using two primers with vector-specific overhangs (SrfAC_o_f and SrfAC_o_r, Table S2). The assembled fragment was cloned into pTrc99a-SrfAC linearized with DraIII and BstBI using InFusion, followed by transformation of Stellar competent cells. After the SOC outgrowth phase, 10 µL of the transformed culture was inoculated in 3 mL of TB medium containing ampicillin and grown overnight. The plasmid library purified from this culture was transformed into HM0079 for protein expression or NEB 5-alpha for long term storage.

***NNK libraries of VSA***

In the VSA gene, 15 positions coding for binding pocket residues were individually targeted for randomization with NNK codons to generate 15 libraries. DNA fragments containing NNK codons were amplified by PCR using pTrc99a-SrfAC-VSA as a template and NNK oligonucleotides as primers. During PCR with NNK oligos, amplification bias might have favoured codons similar to the wild type sequence and thus compromised library quality. To reduce amplification bias, silent mutations adjacent to the NNK codon were added. Depending on the location of the NNK codon in the gene, fragments were generated in one or two steps (Table S3). Where NNK positions were distant from both restriction sites, two fragments were generated and then assembled by PCR using two primers with vector-specific overhangs (VSA_Blp_f and SrfAC_o_r). InFusion cloning was done with one insert fragment carrying the NNK codon and one pTrc99a-SrfAC-VSA vector fragment linearized with appropriate restriction enzymes. Mutants missing from the libraries were cloned and screened in a separate sample batch (Table S5).

### General protein overexpression and purification

For the large-scale expression and purification of individual C-terminally $His_6$-tagged holo-NRPS proteins, a saturated *E. coli* HM0079 (5) culture (0.5 mL) with the appropriate pTrc99a-SrfAC construct was inoculated in 500 mL of 2xYT medium supplemented with ampicillin in a 2 L shaking flask and shaken at 37 °C at 200 rpm. Cultures were grown for 4-6 h until $OD_{600}$ = 1, induced with 0.25 mM isopropyl-D-thiogalactoside (IPTG) and grown for another 16-20 h at 20 °C. Cells were pelleted by centrifugation at 8,000 g and the supernatant was discarded. Cell pellets were resuspended in 30 mL lysis buffer (50 mM TRIS [pH 7.4], 500 mM NaCl, 20 mM imidazole, 2 mM tris(2-carboxyethyl)phosphine [TCEP]). Before cell lysis by sonication, 100 µL of protease inhibitor mix (Sigma, P8849) was added. The lysate was cleared by centrifugation at 19,000 g for 30 min at 4 °C. Proteins were applied to 2 mL of Ni-IDA suspension (Rotigarose, Roth) preequilibrated with lysis buffer by loading the lysate supernatant on the open column. Unbound proteins were removed by washing twice with 20 mL lysis buffer before protein was eluted with 4 x 0.75 mL elution buffer (50 mM TRIS [pH 7.4], 500 mM NaCl, 300 mM imidazole, 2 mM TCEP). Fractions containing protein were pooled and the buffer was exchanged with protein storage buffer (50 mM TRIS [pH 7.6], 200 mM NaCl) on 6 mL Vivaspin (Sartorius) filters with 30 kDa cut-off. Glycerol was added to 10% and the protein concentration adjusted to 50 µM. Samples were flash frozen in liquid nitrogen and stored at -20 °C. To determine the protein concentration, absorbance at 280 nm was measured in Take3 plates on an Epoch2 microplate reader (Biotek) and converted into concentrations using calculated extinction coefficients (www.benchling.com).

##### *SDS-PAGE of overexpressed proteins*

Purity of proteins was determined by SDS-PAGE (Figure S1) using Bolt 4-12% Bis-Tris Plus Gels (ThermoFisher Scientific) with MES-SDS running buffer (Novex). Triple Color Protein Standard III (Serva) was run alongside the protein samples as a size standard. The gels were run at 200 V for 22 min and stained with Quick Coomassie stain (Serva).

#### General hydroxamate specificity assay (HAMA)

##### *Reaction conditions*

The hydroxamate assay (HAMA) for adenylation specificity was conducted with purified proteins as described previously (7). Adenylation reactions of 100 µL contained 50 mM TRIS (pH 7.6), 5 mM $MgCl_2$, 150 mM hydroxylamine (pH 7.5-8, adjusted with NaOH), 5 mM ATP (A2383, Sigma), 1 mM TCEP and 1-5 µM of enzyme. Master mix without the enzyme was prepared and the reaction was initiated by adding enzyme or heat-inactivated enzyme as a control. L-Phe, L-Val and L-Leu were distinguished from D-Phe, D-Val and L-Ile, respectively, by using enantiopure, deuterium labelled standards (EQ Laboratories). Reactions were quenched after 1 h by 10-fold dilution in acetonitrile containing 0.1 % formic acid and immediately analyzed by UPLC-MS/MS. All assays were done from two protein batches in technical triplicates.

##### *UPLC-MS/MS conditions*

Chromatography was performed on a Waters ACQUITY H-class UPLC system (Waters) after injecting 3 µL of the sample. Water with 0.1 % formic acid (A) and acetonitrile with 0.1 % formic acid (B) were used as strong and weak eluent, respectively. Separation of amino acid hydroxamates was achieved on a ACQUITY UPLC BEH Amide column (1.7 µm, 2.1 x 50 mm) with a linear gradient of 10-50% A over 5 min (flow rate 0.4 mL/min) followed by 4 min re-equilibration. Data were analyzed with MassLynx and TargetLynx software (version 4.1). MS/MS detection was performed in positive ion mode on a Xevo TQ-S micro (Waters) tandem quadrupole instrument equipped with an ESI ionisation source. Nitrogen was used as desolvation gas and argon as collision gas. The following source parameters were used: capillary voltage 1.5 kV, cone voltage 65 V, desolvation temperature 500 °C, and desolvation gas flow 1000 L/h. Specific mass transitions recorded in multiple reaction monitoring (MRM) mode were used to detect and quantify amino acid hydroxamates.(7)

### Hydroxamate assay in 96 well plate format

#### *Protein expression*

*E. coli* HM0079 transformed with pTrc99a library constructs was used for overexpression of SrfAC variants in 96 well plate format. Precultures were prepared by inoculating the transformants picked from an agar plate into a round bottom 96-well plate (310 µl, Sarstedt) filled with 150 µl of 2xYT medium supplemented with 100 µg/ml of ampicillin. Each 96-well plate contained four wells with a positive control (pTrc99a-SrfAC for FuncLib library, pTrc99a-SrfAC-VSA for NNK libraries) and 4 wells with a negative control (pTrc99a-SrfAC encoding for SrfAC with disrupted A-domain). Plates were covered with a breathable polyurethane film (Breathe-Easy, Sigma-Aldrich) and incubated for 18 h at 30 °C and 300 rpm in an orbital shaker. The following liquid handling steps were typically performed using a Gilson Platemaster 220 µL as 96-well pipette. For protein expression, 20 µl of the preculture was inoculated into a 96 deep-well plate (2 mL, Sarstedt) containing 1 ml 2xYT medium supplemented with 100 µg/ml ampicillin and incubated for 4-6 h at 30 °C and 300 rpm until the OD600 reached approximately 1. Prior to induction, a 20 µL aliquot was taken from the culture for preparing a 25 % glycerol stock for long-term storage at -80 °C. Additionally, a 5 µL aliquot was taken for sequencing. For induction, the temperature was reduced to 18 °C for 30 min and 0.25 mM IPTG was added (Thermo Scientific). Incubation was continued at 18 °C and 300 rpm for 18-20 h. Cells were harvested by centrifugation at 3000 g and 15 °C and the supernatant was discarded. Immediately before lysis, 50 mL lysis buffer (50 mM TRIS [pH 8.0], 100 mM NaCl, 10 mM imidazole, 1.5 mg/mL lysozyme) were prepared for each plate by adding 50 µL of protease inhibitor mix (P8849, Sigma). The pellet was resuspended in 400 µl lysis buffer per well and the plate was incubated for 30 min at room temperature. Cells were lysed by a single freeze-thaw cycle at -20 °C concluded by thawing the frozen plate for 1.5-2 h at room temperature.

#### *Protein purification*

After thawing, 100 µL of DNA removal mix (50 mM TRIS [pH 8.0], 100 mM NaCl, 10 mM imidazole, 10 mM $MgCl_2$, 10 mM TCEP, 15 U/mL Turbonuclease [Jena Bioscience]) was added to reduce the viscosity of the lysate and incubated without shaking at room temperature for 15 min. Cell debris was removed by centrifugation at 3000 g and 6 °C for 30 min. In a separate, 96-well plate (1.8 mL, Sarstedt) compatible with the magnetic separation rack (S1511S, New England Biolabs), 20 µl of a 25 % Ni-IDA MagBeads (PureCube) suspension was added. The beads were equilibrated with 700 µl lysis buffer and the supernatant was discarded. To purify the released $His_6$-

tagged proteins from the lysate, 400 µl of the lysate supernatant was transferred to the equilibrated beads. The plate was covered with a silicon lid and kept at 4 °C in the fridge for 20 min with vigorous shaking every 5 min to prevent aggregation of the MagBeads. Beads were subsequently pulled down with a magnetic separator and the supernatant was discarded. To remove the unbound proteins and imidazole, the beads were washed twice with 700 µl of wash buffer (50 mM TRIS [pH 8.0], 100 mM NaCl) with the help of the magnetic separator.

##### *Hydroxamate specificity assay (HAMA)*

We found that enzymes maintain adenylation activity on the beads without eluting them. The on-bead assay format was chosen because the imidazole from the elution buffer interferes with subsequent hydroxamate detection. After the second washing step, 100 µl of freshly prepared HAMA master mix (50 mM TRIS [pH 8.0], 5 mM ATP, 5 mM $MgCl_2$, 100 mM hydroxylamine (adjusted to pH 7.5-8 with NaOH), 1 mM TCEP, 1 mM proteinogenic amino acids) was added directly to the beads containing the adsorbed protein and incubated at room temperature for 1.5 h. After the incubation, 6 µl of the reaction mixture was diluted in 54 µl of analysis solution (95% acetonitrile, 0.1% formic acid, 1 µM pipecolic acid hydroxamate as an injection control) in a 384-well plate (100 µL, Brandt). After the dilution step, the 384-well plate was immediately placed on ice and covered with aluminum foil to minimize evaporation of the solvent. The plate was analysed immediately by UPLC-MS/MS according to the general HAMA procedure.

#### Thermostability assay

Thermal stability of SrfAC mutants from the FuncLib library was determined by measuring hydroxamate formation by HAMA after performing reactions for 20 min at different temperatures between 20 and 50 °C. Assays were done with two batches of enzyme in two technical replicates each.

#### Saturation kinetics (MesG/hydroxylamine assay)

Michaelis-Menten parameters of the adenylation with L-Leu for SrfAC and additionally with L-Met and L-Phe for VSA were determined using the MesG/hydroxylamine assay (8). Low activity of SrfAC for L-Phe and L-Met prevented the determination of kinetic parameters. Reactions contained 50 mM TRIS (pH 7.6), 5 mM $MgCl_2$, 100 µM 7-methylthioguanosine (MesG), 150 mM hydroxylamine (adjusted to pH 7.5-8 with NaOH), 5 mM ATP (A2383, Sigma), 1 mM TCEP, 0.4 U/mL inorganic pyrophosphatase (I1643, Sigma), 1 U/mL of purine nucleoside phosphorylase from microorganisms (N8264, Sigma) and 5 µM of enzyme. Flat-bottom 384-well plates (100 µL,

781620, Brand) were used for the reactions. Reactions were started by addition of enzyme and the absorbance was followed at 355 nm on a Synergy H1 (BioTek) microplate reader at 30 °C. Reactions used for background subtraction contained heat-inactivated enzyme. Each substrate concentration was measured in duplicate. Initial velocities (OD $min^{-1}$) were divided by the slope of a pyrophosphate calibration curve to obtain the pyrophosphate release rate. Initial velocities $v_0/[E_0]$ were fit to the Michaelis-Menten equation by nonlinear regression using RStudio version 1.3.1093 (8(9).

#### Data analysis

Random sampling of colonies resulted in a variable number of replicates per mutant. Hydroxamate concentrations were averaged for these replicates within each batch of samples. The total activity of each mutant was calculated as a sum of $N_H = 19$ measured hydroxamates. To minimize the systematic error caused by variable protein expression and purification efficiency between different sample batches, the wild type control measured in each sample batch was used for normalization. A relative activity $A_{rel\,m}$ of each mutant was calculated by dividing the total activity by the total activity of the wild type (Equation 1).

$$A_{rel\,m} = \frac{\sum_{i=1}^{N_H}[\mathrm{HA}]_m}{\sum_{i=19}^{N_H}[\mathrm{HA}]_{wt}} \tag{1}$$

The promiscuity of each mutant was calculated based on the model proposed by Nath et al. (10). The model uses the Shannon entropy $P$ as a metric for promiscuity (Equation 2) with $p_i$ being the probability that the i’th substrate is converted to a hydroxamate by the enzyme.

$$P = \sum_{i=1}^{N_H} p_i \cdot log\, p_i \tag{2}$$

The probability $p_i$ was derived from the proportion of the amino acid hydroxamates (Equation 3).

$$p_i = \frac{[\mathrm{HA}]_i}{\sum_{i=1}^{N_H}[\mathrm{HA}]_\mathrm{i}} \tag{3}$$

Based on $P$, the promiscuity index $I$ of each mutant was calculated as follows:

$$I = -\frac{1}{log\ 19} P \tag{4}$$

*N* indicates the number of measured hydroxamates (N = 19). The promiscuity index $I$ can take values between 0 and 1, with 0 corresponding to perfectly specific and 1 to a perfectly promiscuous enzyme. To better discern the changes in promiscuity caused by mutations, a relative promiscuity index, $I_{rel}$, was calculated by normalization to the wild type (Equation 5). This results in values of $I_{rel}$ larger than 1 for mutants more promiscuous, and lower than 1 for mutants more specific than the wild type.

$$I_{rel} = \frac{I_m}{I_{wt}} \tag{5}$$

Due to the different detection limits of hydroxamates, low adenylation activity results in specificity profiles showing traces of individual products, which results in seemingly low $I$-values. To prevent the false designation of mutants as specific only because promiscuous activities are below the limit of detection, a cut off value for the activity was included. To filter out the mutants showing only traces of activity, before the promiscuity index was calculated, all mutants which accumulate less than 0.2 µM hydroxamates were excluded. Mutants were subsequently ranked according to *I*. *P* and *I* were calculated and visualized in R (Table S6).

#### ***Entropy of activity distribution***

The overall change in activity at one position has been measured as the entropy of the histograms for relative activites (Equation 1) across all 20 mutations in one position. A low entropy value indicates that the overall activity changes a lot when the position is mutated, which usually results from most mutations reducing the activity and only the wild type remaining active.

#### ***Jenson-Shannon distance***

To detect specific mutations that strongly change the substrate profile, we calculated the Jensen-Shannon distance (JSD) between the hydroxamates and the WT production which is bounded between 0 and 1 (11). Given two probability distributions $P$ and $Q$, we calculated the average distribution $R = \frac{1}{2}(P + Q)$ and then

$$JSD(P, Q) = \left(\frac{1}{2}D(P|R) + \frac{1}{2}D(Q|R)\right)^{\frac{1}{2}}, \tag{7}$$

where $D(X|Y)$ denotes the Kullback-Leibler distance between two distributions. The Kullback-Leibler distance was calculated as follows:

$$D(X|Y) = -\sum_i X_i \cdot log\left(\frac{Y_i}{X_i}\right). \quad (8)$$

To avoid that low yielding hydroxamates cause artefacts, it was required that the total activity should be at least 1% of the WT production to be considered for JSD calculation. To determine whether the JSD for a mutation differs significantly from WT, the 95th percentile level of JSD between WT experiments was calculated for WT measured on the same 96-well plate.

##### *Code availability*

The code used for data analysis along with the raw data has been deposited on GitHub:

https://github.com/applied-systems-biology/SrfAC

### Supporting Tables

**Table S1.** Oligonucleotide mix for the FuncLib library of SrfAC.

| Position | Oligo | Molar ratio | Residues | Oligo mix |
|---|---|---|---|---|
| A660 | SrfAC_660_BTT_f | 3 | FLV | SrfAC_660_f |
| | SrfAC_660_GSC_f | 2 | AG | |
| F702 | SrfAC_702_ASC_f | 2 | ST | SrfAC_702_f |
| | SrfAC_702_YAT_f | 2 | HY | |
| | SrfAC_702_TTT_f | 1 | F | |
| | SrfAC_702_TGG_f | 1 | W | |
| C752 | SrfAC_752_TGC_f | 1 | C | SrfAC_752_f |
| | SrfAC_752_DYG_f | 6 | ALMSTV | |

**Table S2.** PCR amplification and the assembly of fragments for FuncLib library of SrfAC.

| Fragment amplification | | Fragment assembly |
|---|---|---|
| **Oligo mix** | **Fragment** | **Oligo** |
| SrfAC_660_f | A | SrfAC_o_f |
| SrfAC_660_r | " | " |
| SrfAC_702_f | B | " |
| SrfAC_702_r | " | SrfAC_o_r |
| SrfAC_752_f | C | " |
| SrfAC_o_r | " | " |

**Table S3.** PCR amplification and the assembly of fragments for NNK libraries of VSA.

| Library | Fragment | Oligo | | Restriction enzyme |
|---|---|---|---|---|
| **VSA-S654NNK** | 654A | VSA_Blp_f | Assembly PCR | BlpI + DraIII |
| | | VSA_S654NNK_o_r | | |
| | 654B | VSA_S654NNK_N655s_f | | |
| | | SrfAC_o_r | | |
| **VSA-F658NNK** | | VSA_F658NNK_D659s_f | | BstBI + DraIII |
| | | SrfAC_o_r | | |
| **VSA-V660NNK** | | VSA_V660NNK_F661s_f | | BstBI + DraIII |
| | | SrfAC_o_r | | |
| **VSA-F661NNK** | | VSA_F661NNK_T662s_f | | BstBI + DraIII |
| | | SrfAC_o_r | | |
| **VSA-F663NNK** | | VSA_F663NNK_D664s_f | | BstBI + DraIII |
| | | SrfAC_o_r | | |
| **VSA-D664NNK** | | VSA_D664NNK_F665s_f | | BstBI + DraIII |
| | | SrfAC_o_r | | |
| **VSA-S702NNK** | 702A | VSA_Blp_f | Assembly PCR | BlpI + DraIII |
| | | VSA_S702NNK_o_r | | |
| | 702B | VSA_S702NNK_A703s_f | | |
| | | SrfAC_o_r | | |
| **VSA-A703NNK** | 703A | VSA_Blp_f | Assembly PCR | BlpI + DraIII |
| | | VSA_A703NNK_o_r | | |
| | 703B | VSA_A703NNK_T704s_f | | |
| | | SrfAC_o_r | | |
| **VSA-L726NNK** | 726A | VSA_Blp_f | Assembly PCR | BlpI + DraIII |
| | | VSA_L726NNK_o_r | | |
| | 726B | VSA_L726NNK_F727s_f | | |
| | | SrfAC_o_r | | |
| **VSA-F727NNK** | 727A | VSA_Blp_f | Assembly PCR | BlpI + DraIII |
| | | VSA_F727NNK_o_r | | |
| | 727B | VSA_F727NNK_G728s_f | | |
| | | SrfAC_o_r | | |
| **VSA-G728NNK** | 728A | VSA_Blp_f | Asse - | BlpI + DraIII |
| | | VSA_G728NNK_o_r | | |

| | | | |
|---|---|---|---|
| | 728B | VSA_G728NNK_G729s_f<br>SrfAC_o_r | |
| **VSA-A752NNK** | | VSA_Blp_f<br>VSA_A752NNK_N751s_r | BlpI + DraIII |
| **VSA-T759NNK** | | VSA_Blp_f<br>VSA_T759NNK_G758s_r | BlpI + DraIII |
| **VSA-V760NNK** | | VSA_Blp_f<br>VSA_V760NNK_T759s_r | BlpI + DraIII |
| **VSA-F761NNK** | | VSA_Blp_f<br>VSA_F761NNK_V760s_r | BlpI + DraIII |

**Table S4**. Oligonucleotide sequences for PCR primers. Targeted positions are labelled in bold.

| Name | Sequence |
|---|---|
| SrfAC_o_f | GATCAGGATACGTTCTTGTCTGTTTC |
| SrfAC_o_r | GAATCCGGCAGATCATGCAC |
| SrfAC_660_BTT_f | GATCAGGATACGTTCTTGTCTGTTTCGAATTACGCCTTTGAT**BTT**TTTACCTTTGATTTCTATGC |
| SrfAC_660_GSC_f | GATCAGGATACGTTCTTGTCTGTTTCGAATTACGCCTTTGAT**GSC**TTTACCTTTGATTTCTATGC |
| SrfAC_660_r | CATGACATTGACATTCTCTTGCAG |
| SrfAC_702_ASC_f | CAAGAGAATGTCAATGTCATG**ASC**GCGACAACCGCACTATTTAATC |
| SrfAC_702_YAT_f | CAAGAGAATGTCAATGTCATG**YAT**GCGACAACCGCACTATTTAATC |
| SrfAC_702_TTT_f | CAAGAGAATGTCAATGTCATG**TTT**GCGACAACCGCACTATTTAATC |
| SrfAC_702_TGG_f | CAAGAGAATGTCAATGTCATG**TGG**GCGACAACCGCACTATTTAATC |
| SrfAC_702_r | GTTAATCAGCTTGCCCGGC |
| SrfAC_752_TGC_f | GCTGCGGATCATGGGGCCGGGCAAGCTGATTAAC**TGC**TACGGGCCGACTGAGGGAAC |
| SrfAC_752_DYG_f | GCTGCGGATCATGGGGCCGGGCAAGCTGATTAAC**DYG**TACGGGCCGACTGAGGGAAC |
| VSA_Blp_f | GATGAAAGAACAAGCGGCTGAGCTG |
| VSA_S654NNK_o_r | ACAGACAAGAACGTATCCTGATCAGAAAATGC |
| VSA_S654NNK_N655s_f | GATACGTTCTTGTCTGTT**NNKAAC**TACGCCTTTGATGTTTTTACCTTTGATTTC |
| VSA_F658NNK_D659s_f | GATCAGGATACGTTCTTGTCTGTTTCGAATTACGCC**NNKGACGTT**TTTACCTTTGATTTCTATGCTTCTATGC |
| VSA_V660NNK_F661s_f | GATCAGGATACGTTCTTGTCTGTTTCGAATTACGCCTTTGAT**NNKTTC**ACCTTTGATTTCTATGCTTCTATGCTG |
| VSA_F661NNK_T662s_f | GATCAGGATACGTTCTTGTCTGTTTCGAATTACGCCTTTGAT**GTTNNKACG**TTTGATTTCTATGCTTCTATGCTGAATGCG |
| VSA_F663NNK_D664s_f | GATCAGGATACGTTCTTGTCTGTTTCGAATTACGCCTTTGATGTTTTTACC**NNKGAC**TTCTATGCTTCTATGCTGAATGCG |
| VSA_D664NNK_F665s_f | GATCAGGATACGTTCTTGTCTGTTTCGAATTACGCCTTTGATGTTTTTACCTTT**NNKTTT**TATGCTTCTATGCTGAATGCGG |
| VSA_S702NNK_o_r | CATGACATTGACATTCTCTTGCAGG |
| VSA_S702NNK_A703s_f | CCTGCAAGAGAATGTCAATGTCATG**NNKGCC**ACAACCGCACTATTTAATCTTCTCAC |
| VSA_A703NNK_o_r | **GCT**CATGACATTGACATTCTCTTGCAGG |
| VSA_A703NNK_T704s_f | CCTGCAAGAGAATGTCAATGTCATG**AGCNNKACC**ACCGCACTATTTAATCTTCTCACAG |
| VSA_L726NNK_o_r | TATACAGCGAAGCCCCTTCATC |
| VSA_L726NNK_F727s_f | GATGAAGGGGCTTCGCTGTATA**NNKTTT**GGCGGAGAGCGCGCGTCAG |
| VSA_F727NNK_o_r | TAATATACAGCGAAGCCCCTTCATC |
| VSA_F727NNK_G728s_f | GATGAAGGGGCTTCGCTGTATATTA**NNKGGT**GGAGAGCGCGCGTCAGTG |
| VSA_G728NNK_o_r | GAATAATATACAGCGAAGCCCCTTC |

**Table S5.** Mutant coverage in NNK libraries of VSA. Numbers in the table denote the frequency of occurrence of the mutant. Missing mutants that were cloned individually with non-degenerate primers are marked in red.

| | Mutation | | | | | | | | | | | | | | | | | | | | |
|---|---|---|---|---|---|---|---|---|---|---|---|---|---|---|---|---|---|---|---|---|---|
| **Position** | **A** | **R** | **N** | **D** | **C** | **Q** | **E** | **G** | **H** | **I** | **L** | **K** | **M** | **F** | **P** | **S** | **T** | **W** | **Y** | **V** | **Missing** |
| **S654** | 3 | 7 | 0 | 2 | 5 | 0 | 3 | 5 | 0 | 4 | 13 | 0 | 2 | 9 | 0 | 10 | 0 | 6 | 3 | 8 | NQHKPT |
| **F658** | 1 | 5 | 3 | 3 | 6 | 3 | 2 | 5 | 0 | 4 | 7 | 3 | 1 | 8 | 2 | 9 | 2 | 5 | 1 | 9 | H |
| **V660** | 1 | 5 | 1 | 3 | 7 | 1 | 1 | 9 | 4 | 6 | 13 | 0 | 1 | 4 | 1 | 4 | 2 | 3 | 4 | 8 | K |
| **F661** | 3 | 3 | 0 | 2 | 0 | 2 | 4 | 10 | 4 | 0 | 5 | 4 | 3 | 7 | 7 | 5 | 1 | 4 | 5 | 11 | NCI |
| **F663** | 1 | 6 | 5 | 2 | 0 | 7 | 2 | 6 | 0 | 1 | 10 | 4 | 3 | 4 | 5 | 1 | 2 | 5 | 0 | 2 | CHY |
| **D664** | 0 | 12 | 5 | 6 | 0 | 4 | 1 | 18 | 0 | 1 | 4 | 0 | 3 | 2 | 4 | 2 | 3 | 0 | 0 | 12 | ACHKWY |
| **S702** | 0 | 4 | 1 | 3 | 1 | 0 | 3 | 8 | 0 | 4 | 8 | 1 | 5 | 10 | 1 | 12 | 0 | 4 | 3 | 7 | AQHT |
| **A703** | 5 | 5 | 0 | 2 | 2 | 0 | 3 | 7 | 4 | 3 | 7 | 1 | 1 | 5 | 1 | 3 | 1 | 7 | 5 | 13 | NQ |
| **L726** | 2 | 4 | 6 | 1 | 5 | 0 | 4 | 4 | 2 | 7 | 7 | 4 | 2 | 8 | 0 | 5 | 2 | 5 | 3 | 6 | QP |
| **F727** | 2 | 1 | 3 | 3 | 6 | 0 | 0 | 5 | 2 | 2 | 14 | 1 | 5 | 11 | 0 | 2 | 3 | 5 | 4 | 7 | QEP |
| **G728** | 2 | 2 | 3 | 2 | 6 | 2 | 2 | 5 | 0 | 4 | 11 | 0 | 5 | 12 | 0 | 3 | 0 | 3 | 3 | 12 | HKPT |
| **A752** | 10 | 6 | 1 | 2 | 4 | 1 | 3 | 1 | 3 | 0 | 11 | 6 | 1 | 3 | 11 | 4 | 4 | 0 | 0 | 1 | IWY |
| **T759** | 1 | 7 | 9 | 0 | 1 | 4 | 1 | 0 | 4 | 5 | 5 | 8 | 0 | 5 | 8 | 8 | 8 | 1 | 5 | 3 | DGM |
| **V760** | 2 | 4 | 2 | 4 | 0 | 2 | 0 | 0 | 4 | 1 | 10 | 10 | 2 | 2 | 6 | 6 | 6 | 1 | 3 | 5 | CEG |
| **F761** | 4 | 2 | 2 | 7 | 1 | 2 | 0 | 2 | 3 | 4 | 7 | 3 | 6 | 5 | 9 | 7 | 4 | 2 | 5 | 5 | E |

**Table S6**. Top 20 mutants from FuncLib SrfAC library and VSA NNK libraries with highest activity ($A_{rel}$), promiscuity and selectivity ($I_{rel}$) relative to the progenitor VSA.

| SrfAC FuncLib library | | | | VSA NNK libraries | | | | | |
|---|---|---|---|---|---|---|---|---|---|
| Activity | | Promiscuity | | Activity | | Promiscuity | | Specificity | |
| Mutant | $A_{rel}$ | Mutant | $I_{rel}$ | Mutant | $A_{rel}$ | Mutant | $I_{rel}$ | Mutant | $I_{rel}$ |
| ASV | 3.35 | VYS | 2.77 | A752I | 2.27 | S702F[#] | 1.46 | G728M | 0.06 |
| ASA | 2.96 | GWV | 2.72 | S702T | 2.21 | V660E | 1.33 | G728L | 0.09 |
| VSA | 2.27 | ASA | 2.66 | V660L | 2.08 | S654I | 1.29 | V660W | 0.10 |
| VSV | 2.03 | GWS | 2.62 | S702A | 2.04 | V660Q | 1.28 | A752M | 0.13 |
| ASL | 1.91 | ASV | 2.45 | V660I | 1.55 | F658Q | 1.27 | V660Y | 0.14 |
| VSL | 1.60 | ASL | 2.34 | A703N | 1.54 | V660S | 1.27 | V660F | 0.21 |
| LSL | 1.51 | AWS | 2.31 | V660A | 1.51 | V660A | 1.27 | F761A | 0.27 |
| GSL | 1.46 | AWM | 2.28 | A703I | 1.51 | F658A | 1.26 | G728A | 0.29 |
| VFA | 1.39 | VSA | 2.24 | F663W | 1.51 | S654Q | 1.26 | F727Y | 0.41 |
| GTV | 1.28 | VFA | 2.19 | A703M | 1.42 | F658S | 1.25 | S702D | 0.41 |
| GSV | 1.22 | GWC | 2.14 | F661A | 1.41 | F663F | 1.24 | L726D | 0.48 |
| GTL | 1.15 | FWL | 2.00 | S654N | 1.37 | F658G | 1.24 | L726G | 0.50 |
| GSC | 1.07 | FFA | 1.98 | V760G | 1.33 | A752G | 1.22 | F727S | 0.50 |
| GST | 1.05 | VSV | 1.91 | A703L | 1.24 | F658T | 1.21 | G728F | 0.51 |
| VFM | 1.03 | LWA | 1.86 | A752V | 1.24 | S654L | 1.21 | L726A | 0.57 |
| AFC | 1.02 | GYM | 1.69 | S654A | 1.23 | D664E | 1.20 | F761I | 0.57 |
| FSV | 0.99 | FWS | 1.66 | S654G | 1.19 | S654G | 1.20 | F727A | 0.57 |
| GYV | 0.98 | AWL | 1.63 | F727I | 1.15 | F661T | 1.20 | F761V | 0.58 |
| FSA | 0.96 | VYL | 1.63 | A703A | 1.10 | S654M | 1.20 | L726Y | 0.58 |
| FSL | 0.96 | GSL | 1.54 | V660G | 1.07 | F658H | 1.19 | F727T | 0.58 |

[#]The high promiscuity of this mutant could not be confirmed with protein purified on large scale.

**Table S7**. Specificity codes of SrfAC and mutants compared to natural specificity codes. According to the measured specificity, mutants have been classified into A-domain types. For natural A-domains of the same type according to a database of specificities and codes (12), the frequency of occurrence of residues at the specified positions is noted as percentage. Residue identities that are absent (0 %), rare (0 - 10 %), or frequent (>10 %) are marked in red, yellow, and light green, respectively.

| | | Specificity code position in SrfAC numbering<br>Second row: percentage in natural codes | | | | | | | | | |
|---|---|---|---|---|---|---|---|---|---|---|---|
| **Enzyme** | **Measured specificity** | **659** | **660** | **663** | **702** | **726** | **728** | **752** | **760** | **761** | **A-domain type according to Rausch et al.** (12) |
| **GrsA** | Phe | D | A | W | T | I | A | A | I | C | |
| **SrfAC** | Leu | D | A | F | F | L | G | C | V | F | |
| **VSA** | multi | D | V<br>2% | F | S<br>22% | L | G | A<br>83% | V | F | nonpolar |
| **Met-specific** | Met | D | M/I/<br>L | F | S | L | G | A | V | F | - |
| **A752G** | Phe | D | V<br>0% | F | S<br>0% | L | G | G<br>23% | V | F | Phe |
| **V660A** | aromatic | D | A | F | S<br>7% | L | G | A<br>64% | V | F | aromatic |
| **S702A** | aromatic | D | V<br>1.5% | F | A<br>6% | L | G | A<br>64% | V | F | aromatic |
| **V760G** | D/L-Phe | D | V<br>0% | F | S<br>0% | L | G | A<br>77% | G<br>0% | F | Phe |
| **G728A** | Ala | D | V<br>3% | F | S<br>0% | L | A<br>37% | A<br>10% | V | F | small |
| **G728M** | Ala | D | V<br>3% | F | S<br>0% | L | M<br>0% | A<br>10% | V | F | small |
| **V660W** | Leu | D | W<br>0% | F | S<br>0% | L | G | A<br>10% | V | F | Leu |
| **V660F** | Leu | D | F<br>1% | F | S<br>0% | L | G | A<br>10% | V | F | Leu |

#### Supporting Figures

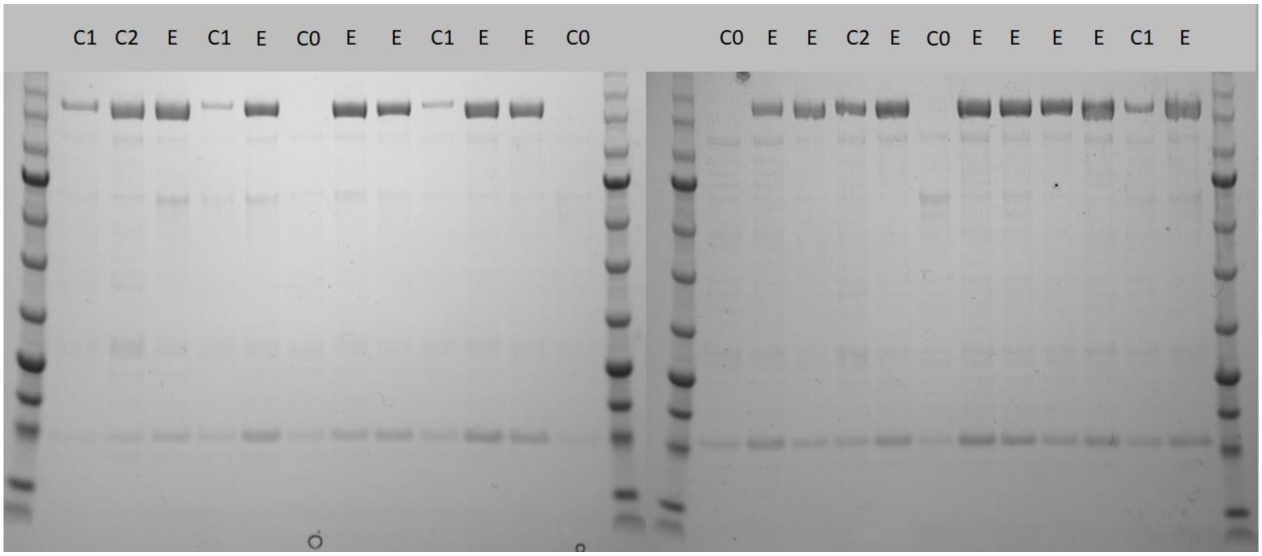

**Figure S1.** SDS PAGE of SrfAC expressed and purified in 96-well plate format. Proteins were eluted from magnetic beads with 200 µL of elution buffer (50 mM TRIS pH 8.0, 200 mM imidazole) and 5 µL was loaded on the gel. E, HM0079 strain with pTrc99a-SrfAC; C0, negative control containing the empty vector; C1, purification control with empty vector and SrfAC added to the cell lysate; C2, purification control with empty vector and SrfAC added to the eluate.

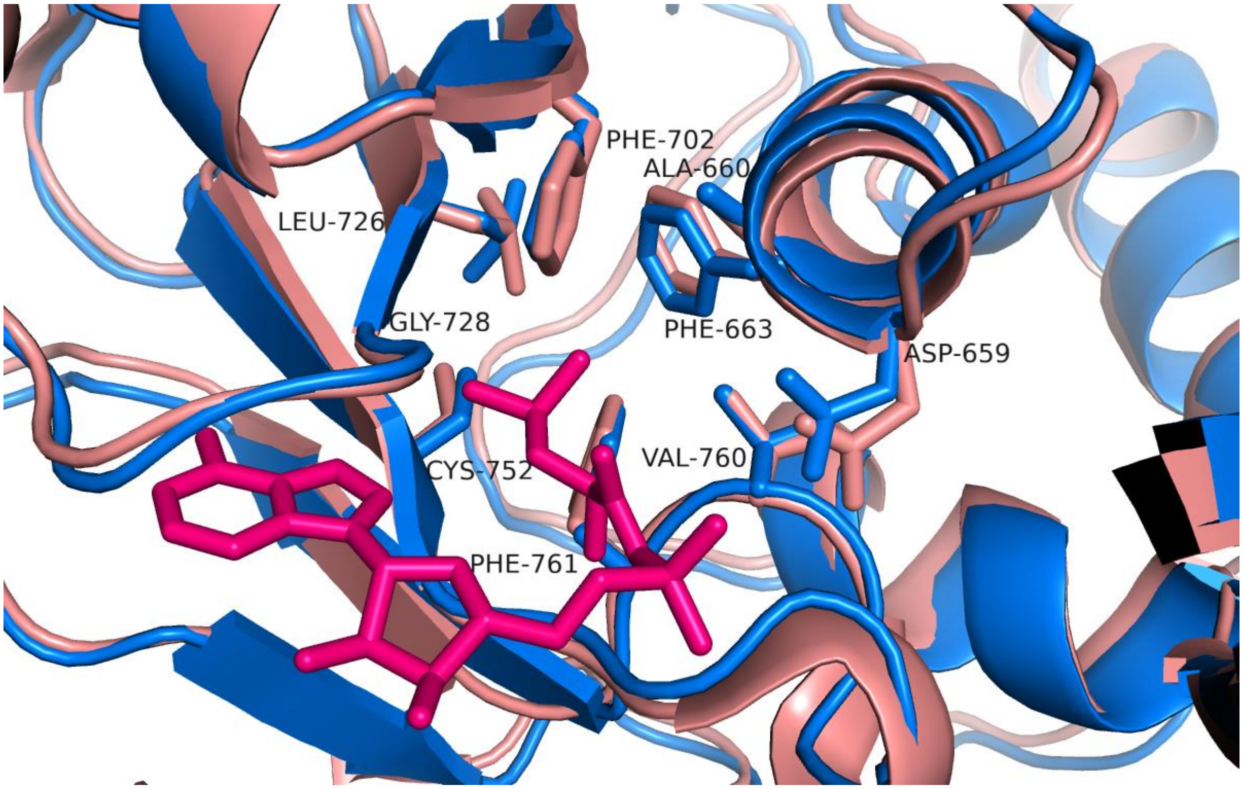

**Figure S2.** Overlay of YASARA model of SrfAC with Leu-AMP (blue) and SrfAC crystal structure (PDB: 2VSQ, pink). Specificity code residues are labeled.

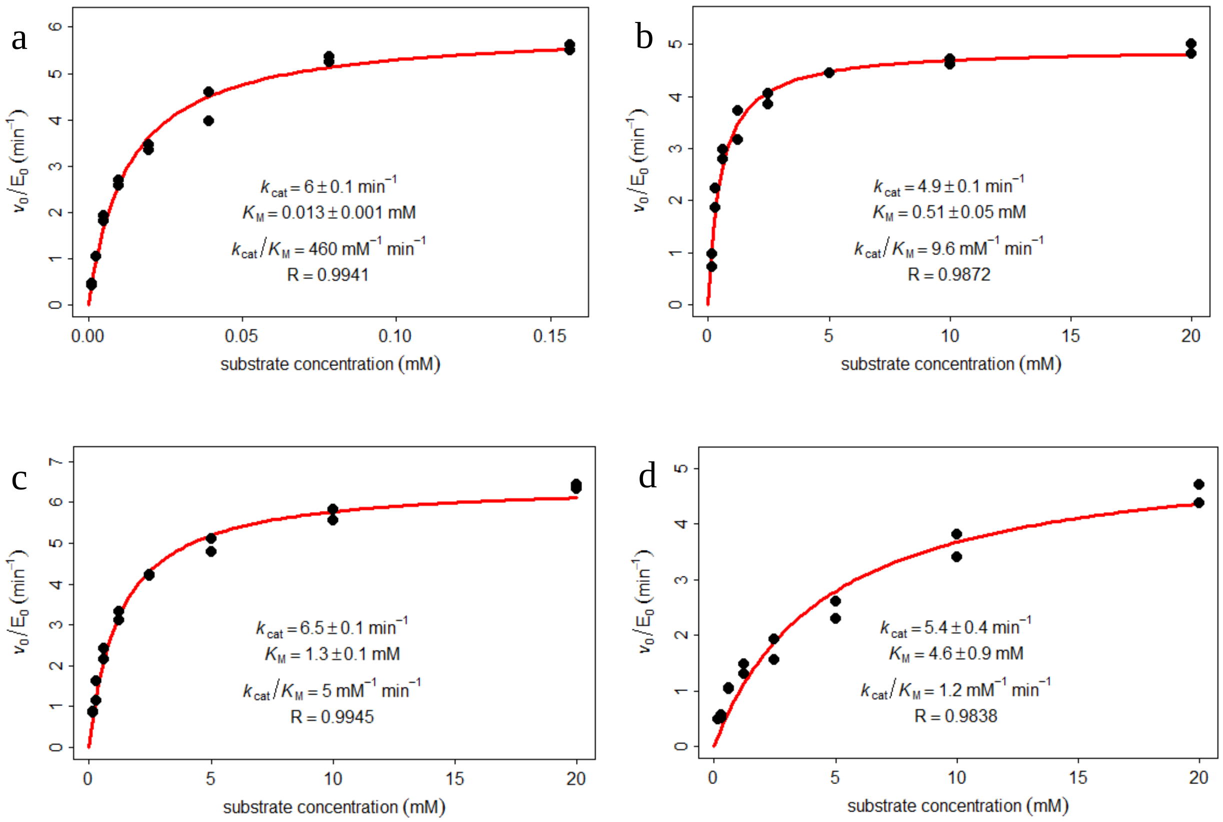

**Figure S3**. Saturation kinetics of SrfAC with L-Leu (a) and VSA with L-Leu (b), L-Phe (c) and L-Met (d) measured with the MesG/hydroxylamine spectrophotometric assay. Assays were conducted with a single enzyme batch in technical duplicates.

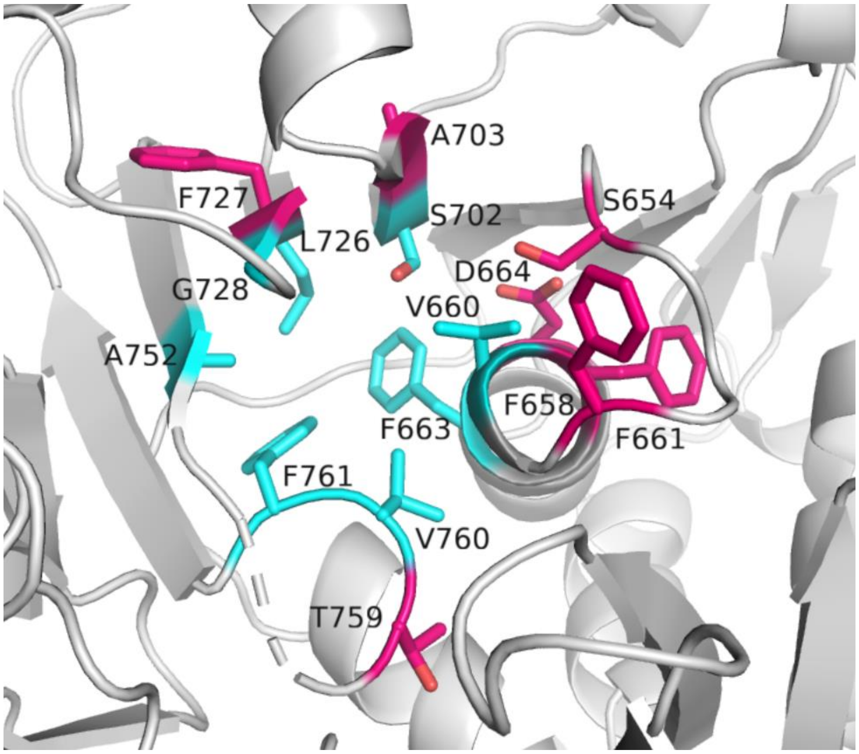

**Figure S4**. Residues in the binding pocket of VSA model selected for saturation mutagenesis. The specificity code residues in the first shell are shown in cyan and second shell in pink. The VSA structure is SWISS homology model built against SrfAC (PDB: 2vsq) as a template.

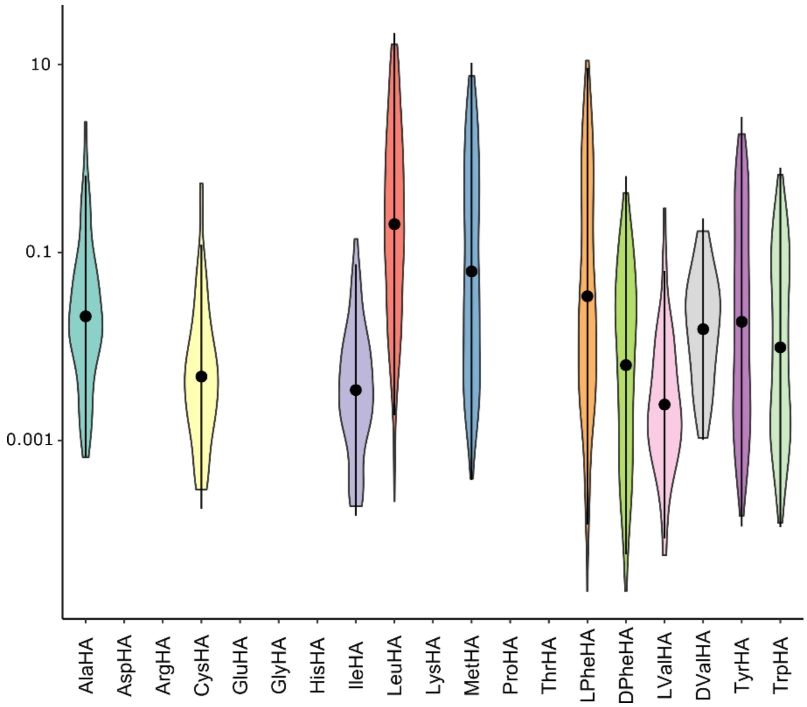

**Figure S5**. Logarithmic distribution of concentration (μM) of detected hydroxamates pooled from 15 NNK libraries.

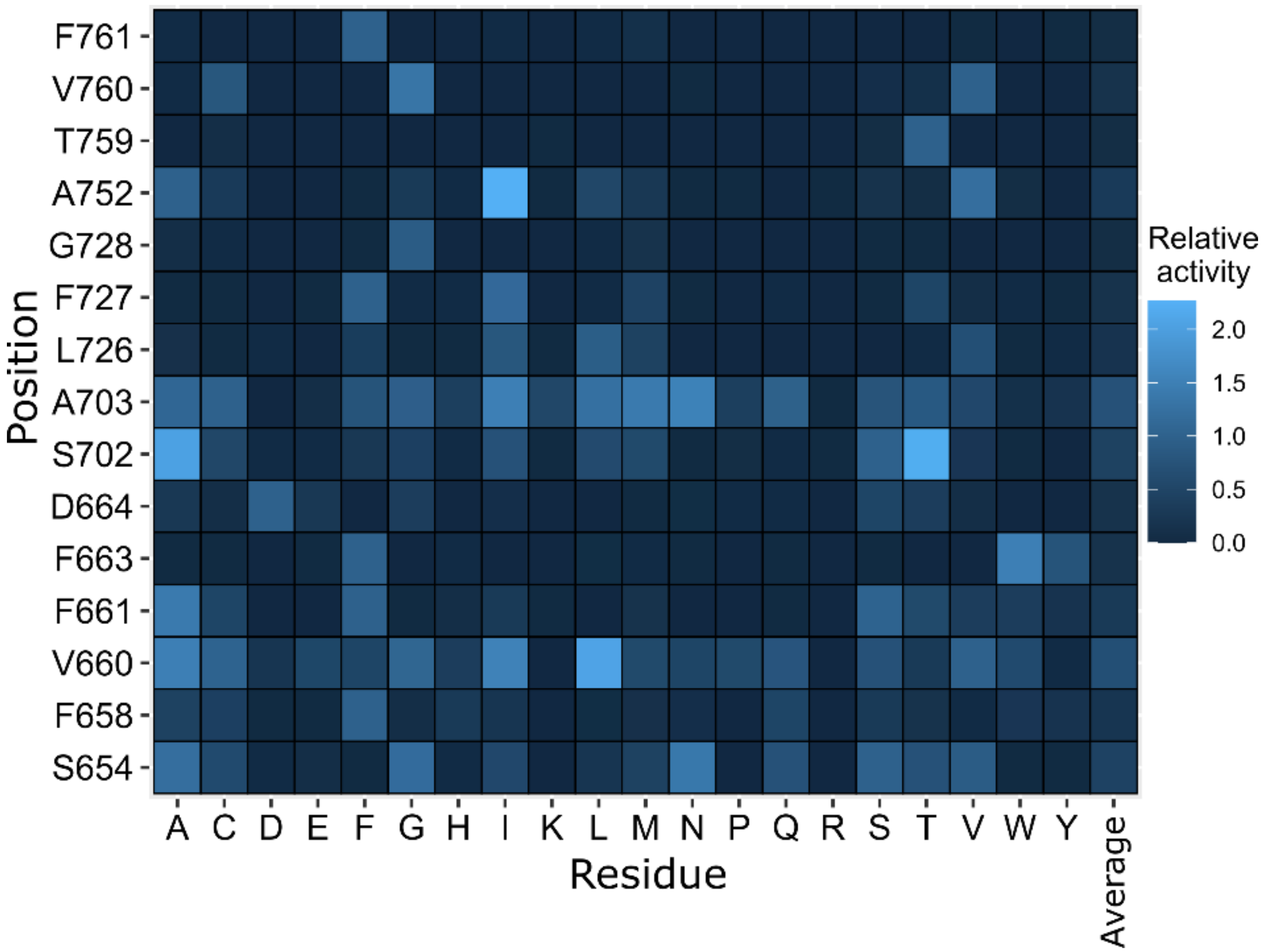

**Figure S6**. Heatmap of activities of all mutants from 15 NNK libraries relative to the progenitor VSA. Activity is calculated as a sum of all formed hydroxamates per mutant. Last column represents the average activity per position.

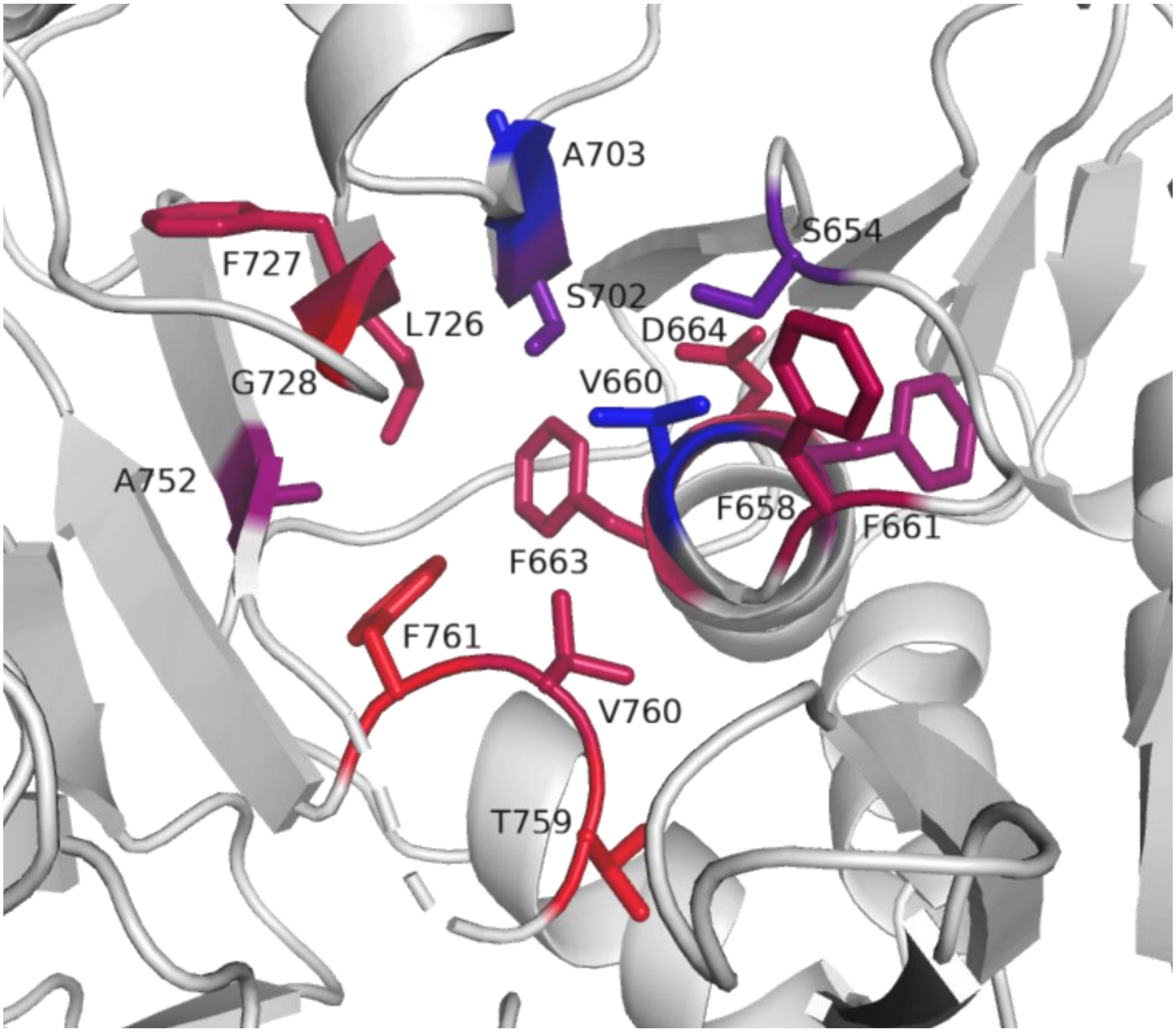

**Figure S7**. Binding pocket of VSA homology model with targeted residues colored according to the average activity per position, relative to the progenitor VSA. Mutations at blue positions result in highest activities and at red positions, lowest.

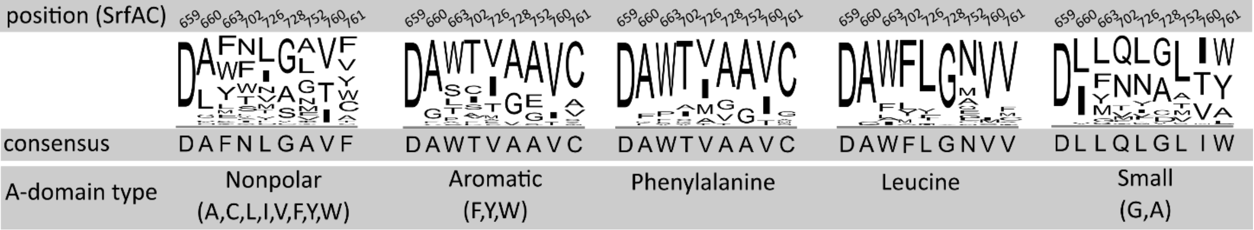

**Figure S8**. Consensus sequences of specificity codes of A-domains activating different amino acid substrates. Alignments are generated with Muscle (13) using curated specificity code database from Rausch et al. (12).

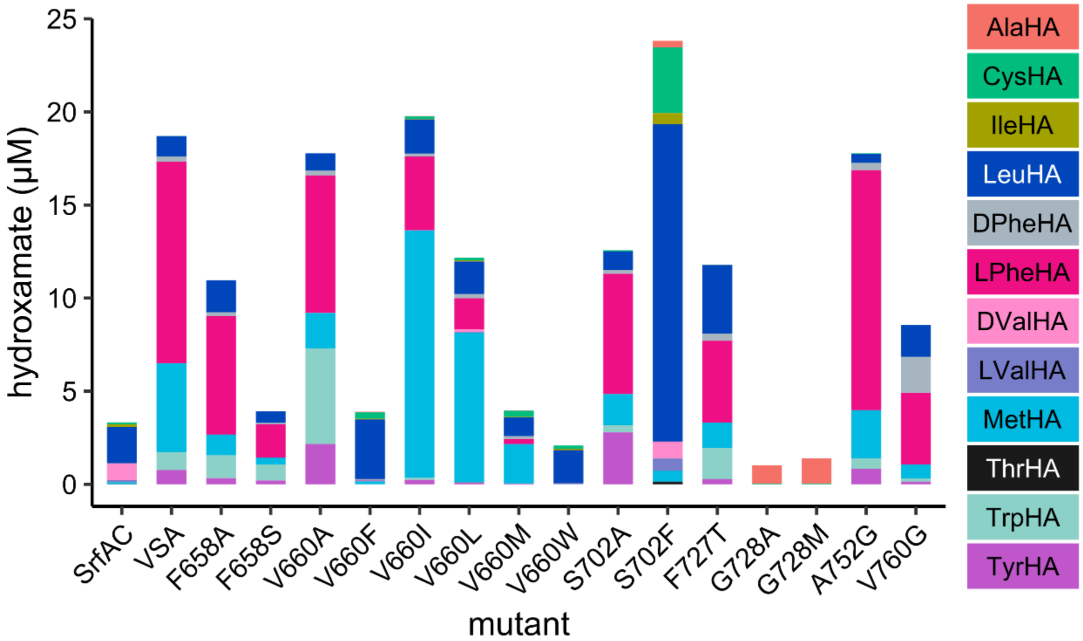

**Figure S9**. HAMA profiles of SrfAC, VSA, and mutants. Fractions of hydroxamates are means from three technical replicates from two batches of enzyme. Enzyme reactions were incubated for 60 min at 25 ºC and 1 µM enzyme. The plot is a different representation of the data shown in Figure 4.

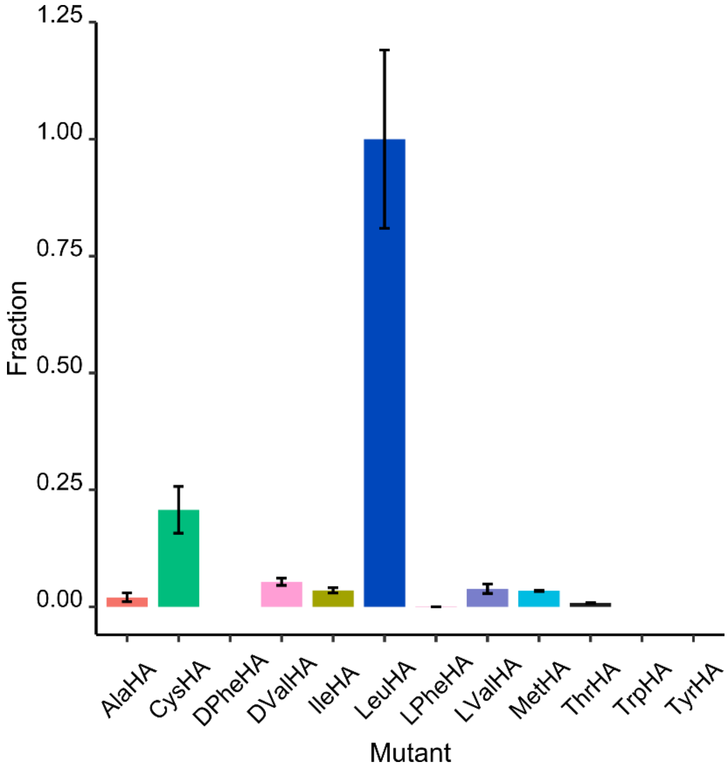

**Figure S10**. HAMA profile of S702F mutant of VSA. Error bars are standard deviations from three technical replicates from two batches of enzyme. Enzyme reactions were incubated for 60 min at 25 ºC and 1 µM enzyme.

### Sequences of proteins used in this study

The A-domain is highlighted in blue. The residues mutated in the VSA mutant are highlighted in bold, randomized residues in the specificity code in red, and those in the second shell in yellow.

#### SrfAC

MSQFSKDQVQDMYYLSPMQEGMLFHAILNPGQSFYLEQITMKVKGSLNIKCLEESMNVIMDRYDVFRTVFIHEKVKRPVQVVLKKRQ
FHIEEIDLTHLTGSEQTAKINEYKEQDKIRGFDLTRDIPMRAAIFKKAEESFEWVWSYHHIILDGWCFGIVVQDLFKVYNALREQKP
YSLPPVKPYKDYIKWLEKQDKQASLRYWREYLEGFEGQTTFAEQRKKQKDGYEPKELLFSLSEAETKAFTELAKSQHTTLSTALQAV
WSVLISRYQQSGDLAFGTVVSGRPAEIKGVEHMVGLFINVVPRRVKLSEGITFNGLLKRLQEQSLQSEPHQYVPLYDIQSQADQPKL
IDHIIVFENYPLQDAKNEESSENGFDMVDVHVFEKSNYDLNLMASPGDEMLIKLAYNENVFDEAFILRLKSQLLTAIQQLIQNPDQP
VSTINLVDDREREFLLTGLNPPAQAHETKPLTYWFKEAVNANPDAPALTYSGQTLSYRELDEEANRIARRLQKHGAGKGSVVALYTK
RSLELVIGILGVLKAGAAYLPVDPKLPEDRISYMLADSAAACLLTHQEMKEQAAELPYTGTTLFIDDQTRFEEQASDPATAIDPNDP
AYIMYTSGTTGKPKGNITTHANIQGLVKHVDYMAFSDQDTFLSVSNYAFDAFTFDFYASMLNAARLIIADEHTLLDTERLTDLILQE
NVNVMFATTALFNLLTDAGEDWMKGLRCILFGGERASVPHVRKALRIMGPGKLINCYGPTEGTVFATAHVVHDLPDSISSLPIGKPI
SNASVYILNEQSQLQPFGAVGELCISGMGVSKGYVNRADLTKEKFIENPFKPGETLYRTGDLARWLPDGTIEYAGRIDDQVKIRGHR
IELEEIEKQLQEYPGVKDAVVVADRHESGDASINAYLVNRTQLSAEDVKAHLKKQLPAYMVPQTFTFLDELPLTTNGKVNKRLLPKP
DQDQLAEEWIGPRNEMEETIAQIWSEVLGRKQIGIHDDFFALGGHSLKAMTAASRIKKELGIDLPVKLLFEAPTIAGISAYLKNGGS
DGLQDVTIMNQDQEQIIFAFPPVLGYGLMYQNLSSRLPSYKLCAFDFIEEEDRLDRYADLIQKLQPEGPLTLFGYSAGCSLAFEAAK
KLEEQGRIVQRIIMVDSYKKQGVSDLDGRTVESDVEALMNVNRDNEALNSEAVKHGLKQKTHAFYSYYVNLISTGQVKADIDLLTSG
ADFDMPEWLASWEEATTGVYRVKRGFGTHAEMLQGETLDRNAEILLEFLNTQTVTVS

#### SrfAC-VSA

MSQFSKDQVQDMYYLSPMQEGMLFHAILNPGQSFYLEQITMKVKGSLNIKCLEESMNVIMDRYDVFRTVFIHEKVKRPVQVVLKKRQ
FHIEEIDLTHLTGSEQTAKINEYKEQDKIRGFDLTRDIPMRAAIFKKAEESFEWVWSYHHIILDGWCFGIVVQDLFKVYNALREQKP
YSLPPVKPYKDYIKWLEKQDKQASLRYWREYLEGFEGQTTFAEQRKKQKDGYEPKELLFSLSEAETKAFTELAKSQHTTLSTALQAV
WSVLISRYQQSGDLAFGTVVSGRPAEIKGVEHMVGLFINVVPRRVKLSEGITFNGLLKRLQEQSLQSEPHQYVPLYDIQSQADQPKL
IDHIIVFENYPLQDAKNEESSENGFDMVDVHVFEKSNYDLNLMASPGDEMLIKLAYNENVFDEAFILRLKSQLLTAIQQLIQNPDQP
VSTINLVDDREREFLLTGLNPPAQAHETKPLTYWFKEAVNANPDAPALTYSGQTLSYRELDEEANRIARRLQKHGAGKGSVVALYTK
RSLELVIGILGVLKAGAAYLPVDPKLPEDRISYMLADSAAACLLTHQEMKEQAAELPYTGTTLFIDDQTRFEEQASDPATAIDPNDP
AYIMYTSGTTGKPKGNITTHANIQGLVKHVDYMAFSDQDTFLSVSNYAFD**V**FTFDFYASMLNAARLIIADEHTLLDTERLTDLILQE
NVNVM**S**ATTALFNLLTDAGEDWMKGLRCILFGGERASVPHVRKALRIMGPGKLIN**A**YGPTEGTVFATAHVVHDLPDSISSLPIGKPI
SNASVYILNEQSQLQPFGAVGELCISGMGVSKGYVNRADLTKEKFIENPFKPGETLYRTGDLARWLPDGTIEYAGRIDDQVKIRGHR
IELEEIEKQLQEYPGVKDAVVVADRHESGDASINAYLVNRTQLSAEDVKAHLKKQLPAYMVPQTFTFLDELPLTTNGKVNKRLLPKP
DQDQLAEEWIGPRNEMEETIAQIWSEVLGRKQIGIHDDFFALGGHSLKAMTAASRIKKELGIDLPVKLLFEAPTIAGISAYLKNGGS
DGLQDVTIMNQDQEQIIFAFPPVLGYGLMYQNLSSRLPSYKLCAFDFIEEEDRLDRYADLIQKLQPEGPLTLFGYSAGCSLAFEAAK
KLEEQGRIVQRIIMVDSYKKQGVSDLDGRTVESDVEALMNVNRDNEALNSEAVKHGLKQKTHAFYSYYVNLISTGQVKADIDLLTSG
ADFDMPEWLASWEEATTGVYRVKRGFGTHAEMLQGETLDRNAEILLEFLNTQTVTVS
